# PE11 promotes intracellular persistence of *Mycobacterium tuberculosis* by inhibiting autophagy and lysosomal biogenesis by targeting the FLCN-lactate-TFEB signaling axis

**DOI:** 10.1101/2025.07.31.668032

**Authors:** Priyanka Dahiya, Manoj Kumar Bisht, Abhishek Saha, Aakash Chandramouli, Siddhesh S. Kamat, Vinay Kumar Nandikoori, Sudip Ghosh, Sangita Mukhopadhyay

**Author notes:** To whom correspondence should be addressed: Laboratory of Molecular Cell Biology, BRIC-CDFD, Hyderabad - 500039, Telangana, India.

## Abstract

*Mycobacterium tuberculosis* (Mtb) employs multiple virulence factors, including cell wall-associated proteins, to evade host immune responses. PE11, a cell wall-localized esterase, contributes to Mtb persistence by facilitating cell wall remodelling and resistance to acidic and antibiotic stress. Herein we describe a novel role of PE11 in subverting host autophagy through disruption of TFEB-mediated lysosomal function. PE11 promotes FLCN-dependent depletion of intracellular lactate to destabilize TFEB and thereby downregulate genes essential for autophagic flux and lysosomal acidification. Using a PE11-deficient Mtb strain, we demonstrate that PE11 targets the FLCN–lactate axis to regulate TFEB stability. Exogenous lactate supplementation restored TFEB stability, enhanced lysosomal acidification, and significantly reduced intracellular bacterial burden. Lactate also synergized with frontline anti-tubercular drugs to improve Mtb clearance. These findings establish PE11 as a key immune evasion factor and highlight lactate as a promising host-directed therapeutic to enhance bacterial killing and reduce antibiotic-associated toxicity.

## Introduction

Tuberculosis (TB), caused by *Mycobacterium tuberculosis* (Mtb), remains a global health crisis, claiming about 1.25 million lives annually (Global Tuberculosis Report, 2024). A major challenge in controlling TB is the bacterium’s ability to persist in its latent form (LTBI) in an estimated quarter of the global population (Lillebaek et al., 2002; Houben et al., 2016). During this latency, the infection stays asymptomatic as Mtb uses host lipids to meet its energy needs, keeping itself in a dormant or growth-arrested state. These bacilli often hide in protective areas like granulomas, adipose tissue (Neyrolles et al., 2006; Ayyappan et al., 2019), and mesenchymal stem cells (Jain et al., 2019).

The success of Mtb as one of the deadliest pathogens to the humankind is due to presence of an extremely resilient cell wall and adoption of several strategies to evade the host immune system. The cell wall significantly contributes to its ability to resist various stressors, such as acidic, oxidative stress, and several common antibiotics in use (Velayati et al., 2009; Kalscheuer et al., 2019). This complex structure, composed mainly of peptidoglycan, arabinogalactan and mycolic acid which provides a hydrophobic barrier that protects the bacterium from the host’s immune system (Brennan and Nikaido, 1995). The Mtb genome encodes for several lipolytic enzymes which play crucial roles in the lipid and fatty acid metabolism, cell wall remodelling as well as virulence of the bacteria (Lin et al., 2024). Some of these enzymes are involved in disrupting phagosome functions by degrading the phagosomal membrane and modifying its permeability while some others are known to possess immunomodulatory properties (Lin et al., 2024).

Earlier, we have shown that a cell wall-localized protein PE11 (also known as LipX, Rv119c), is an esterase that can significantly remodel the mycobacterial cell wall (Singh et al., 2016). PE11, is the smallest member (∼10.9 kDa) of the Lip family, is also is a member of the PE family of Mtb (Dahiya et al., 2024). PE11 is present in pathogenic strains like Mtb, *M. bovis*, and the clinical strain Mtb like CDC1551, but conspicuously absent in the non-pathogenic strain like *M. smegmatis*. Expression of PE11 was found to be markedly upregulated in response to various stressors like H_2_O_2_, SDS, and Diamide (Voskuil et al., 2011; Fontán et al., 2009) as well as during starvation and in hypoxic lipid-loaded macrophages (Betts et al., 2002; Voskuil et al, 2004; Daniel et al, 2011). It is also found to be upregulated during infection in macrophages as well as in granulomas of the lung in active TB patients (Schnappinger et al., 2003; Sampson, 2011; Rachman et al., 2006). In addition, PE11 has been shown to play a role in inducing necrotic death of infected macrophages (Deng et al., 2015). Expression of PE11 in *M. smegmatis* was found to enhance cell surface hydrophobicity and alter colony morphology, biofilm formation, and antibiotic susceptibility (Singh et al., 2016). In addition, PE11 was found to modulate production of pro-inflammatory cytokines (Singh et al., 2016; Deng et al., 2015).

Herein, we uncover a novel function of PE11 in addition to its cell wall remodelling and inflammatory functions. Using a *pe11* knock-out strain of *M. tuberculosis* (H37Rv), coupled with transcriptomic and functional studies, we demonstrate that PE11 plays a pivotal role in immune evasion by interfering with autophagy-lysosome function. The autophagy-lysosome system is crucial for clearing intracellular pathogens by transporting them to lysosomes for degradation (Kaufmann et al., 2018; Sapkota et al., 2024). PE11 was found to downregulate key autophagy-associated genes as well as TFEB, a master transcriptional regulator for lysosomal gene expression (Settembre et al., 2011; Napolitano & Ballabio, 2016). Mechanistically, PE11 was found to affect the stability of TFEB, by reducing cellular lactate levels through regulation of a tumour suppressor gene, Folliculin (FLCN). This FLCN-lactate-TFEB axis appears to be a novel immune-subversive mechanism exploited by Mtb through PE11. In addition, exogenous lactate supplementation was found to restore TFEB stability and autophagic function, reducing mycobacterial burden *in vitro* and *in vivo*, and synergizing with first-line antibiotics. Thus, we demonstrated lactate to be a potential candidate for development of host-directed therapy to control TB.

## Material and Methods

### Antibodies and reagents

The detail description of antibodies and reagents used in this study is described in Supplementary Table 1

### Mammalian cell culture

BMC2 macrophages were cultured in Dulbecco’s Modified Eagle Medium (DMEM, Hyclone, USA) supplemented with 10% Fetal bovine serum (FBS, Gibco, USA) and 1X Antibiotic-Antimycotic solution (Gibco, USA) and 1% Glutamax (Invitrogen, ThermoFisher Scientific, USA) [DMEM-10].

### Mice

C57BL/6 mice were maintained at the animal house facility of the BRIC-Centre for DNA Fingerprinting and Diagnostics (CDFD), Hyderabad. Experiments were approved by the Institutional Animal Ethics Committee (IAEC). All experiments were performed as per the guidelines of the IAEC of BRIC-CDFD.

### Isolation of mouse peritoneal macrophages

About 8 weeks old C57BL/6 mice were given one ml intraperitoneal injection of 4% thioglycolate medium. Three days post-injection, mice were sacrificed using CO_2_ asphyxiation and peritoneal macrophages were harvested as described earlier (Bisht et al., 2023). The adherent macrophages were cultured in DMEM-10.

### Generation and culture of mycobacterial strains

Mycobacterial strains like *M. smegmatis* and *M. tuberculosis* H37Rv were cultured in Middlebrook 7H9 medium (BD DifcoTM, USA) supplemented with 10% albumin, dextrose, NaCl, catalase (ADC) along with 0.2% glycerol (Sigma-Aldrich, USA), and 0.05% Tween-80 (Sigma-Aldrich, USA) or plated in 7H10/7H11 agar (BD DifcoTM, USA) with 10% OADC (HIMEDIA, India) and 0.2% glycerol. *M. smegmatis* strain expressing PE11 (*Msmeg*-PE11) or harbouring the backbone vector pVV16 (*Msmeg*-pVV) and *M. tuberculosis* PE11 knock-out H37Rv strain (H37Rv-*pe11*KO) were generated as described earlier (Dahiya et al., 2024). The H37Rv-*pe11*KO strain was created using phage-mediated recombineering, with ∼850 bp (LHS) and ∼950 bp (RHS) flanking regions amplified, ligated with HygR, and cloned into phAE159. Complemented strain (H37Rv-*pe11*KO::*pe11*) was generated using a theophylline-inducible riboswitch vector with wild-type *pe11* electroporated into H37Rv-*pe11*KO.

### Scanning electron microscopy (SEM)

For scanning electron microscopy of *M. tuberculosis* bacteria, cells were harvested by low-speed centrifugation, washed in sterile phosphate-buffered saline (PBS), and fixed for a minimum of 2 hours in 4% (vol/vol) formaldehyde in PBS. Cells were then pelleted, supernatant was removed and washed with 0.1 M Sodium cacodylate buffer, and the pellet was resuspended in 1% (wt/vol) osmium tetroxide for 3 hours. Cells were then dehydrated through a graded ethanol series (50, 70, 90, and 100%; each level was applied for 10 minutes each time). Carbon and silver tape were coated followed by a smear of cells. Further gold particles are coated onto the bacterial cells and scanning electron microscopy was performed.

### Infection of macrophages

BMC2 macrophages or peritoneal macrophages from C57BL/6 mice were infected with various *M. smegmatis* strains at 1:10 MOI (multiplicity of infection) for 3 hours followed by washing to remove the extracellular bacteria and incubated for another 3 hours. Cells were then harvested for carrying out various experiments. Also, BMC2 or C57BL/6 peritoneal macrophages were infected with various *M. tuberculosis* strains at 1:5 MOI for 4 hours followed by washing to remove the extracellular bacteria and further incubated for another 12 hours. Culture supernatants were harvested for estimation of various cytokines (IL-1β, IL-6, TNF-α) and cells were harvested for various experiments. Infection with Mtb was carried out in the BSL3 facility of Centre for Cellular and Molecular Biology (CCMB), Hyderabad, India.

### Infection of mice

C57BL/6 mice were intraperitoneally infected with 50 million bacilli of either *Msmeg*-pVV or *Msmeg*-PE11 and blood as well as lung tissues were collected three days post-infection for various experiments. Also, mice were exposed to approx. 20 CFU *M. tuberculosis* strain H37Rv (wild-type) or PE11 knock-out strain (H37Rv-*pe11*KO) or complemented strain containing wild-type PE11 (H37Rv-*pe11*KO::*pe11*) through aerosol route at the ABSL3 facility of the ‘International Centre for Genetic Engineering and Biotechnology (ICGEB), New Delhi, India’. To confirm bacterial deposition, two mice per group were sacrificed 24 hours post-infection. After 4 weeks, mice were euthanized. Blood was collected for cytokine analysis from retro-orbital route. Also, lungs were collected for bacterial burden as well as for carrying out histopathology and real-time PCR (RT-PCR) of various genes.

### Histopathology

For histopathology, 10% paraformaldehyde-fixed lung tissues were embedded in paraffin and cut into 4.5 μm thick sections. Replicate sections were stained with Hematoxylin and Eosin (H&E) to assess granuloma formation and inflammation.

### Blood profiling

Blood was collected from retro-orbital route in 4% EDTA tubes from C57BL/6 mice infected with various *M. smegmatis* strains. For complete blood profile, blood samples were analyzed on automatic cell analyzers, ADVIA 2120 (Hematology System; Siemens Healthcare Diagnostics, Forchheim, Germany).

### Confocal microscopy

Confocal microscopy was carried out for studying LC3B and TFEB proteins. In brief, BMC2 macrophages were infected with various mycobacterial strains, fixed with 4% paraformaldehyde and permeabilized with 0.05% Triton-X, and probed with specific combinations of primary and secondary antibodies (Abs). Cells were mounted with DAPI-containing Vectashield mounting medium (Vector Laboratories, USA) and visualized using Leica Microsystem Microscope (Germany).

### Western blotting

Western blotting was carried out to check the expression levels of p62, LC3B, FLCN and β-actin in macrophages infected with various mycobacterial strains. For this, protein samples were extracted from C57BL/6 peritoneal macrophages infected with various mycobacterial strains using the lysis buffer containing NP-40 and subjected to centrifugation to remove cellular debris. The soluble protein fraction was separated based on molecular weight by sodium dodecyl sulfate-polyacrylamide gel electrophoresis (SDS-PAGE) and transferred onto a polyvinylidene fluoride (PVDF) membrane. The membrane was incubated with specific combinations of primary and secondary antibodies, followed by chemiluminescent detection using a commercially available kit (GE Healthcare, UK) according to the manufacturer’s instructions.

### Enzyme-linked immunosorbent assay (ELISA)

The levels of IL-1β, IL-6, and TNF-α were determined in both blood sera, cell culture supernatants and lung extracts using a commercially available mouse ELISA kit (Invitrogen, ThermoFisher Scientific, USA). The assay was performed according to the manufacturer’s protocol, and absolute concentration of IL-1β/IL-6/TNF[α cytokine was measured using a standard curve provided by the manufacturer.

### Bacterial survival assay

For CFU count, lung tissues were homogenized in saline solution and serial dilutions of lysates were plated on 7H11 plates supplemented with 10% OADC (HIMEDIA, India). Plates were incubated at 37°C and colonies were counted after 3-4 weeks. Bacillary loads in the lungs of infected mice were evaluated 4 weeks post-infection. In another study, thioglycolate-elicited peritoneal macrophages (0.5 × 10^6^) from C57BL/6 mice were infected with various *M. tuberculosis* H37Rv strains at 1:5 MOI for 4 hours in the absence or presence of lactate (10 mM and 20 mM) either as standalone or in combination with Rifampicin (1 µg/ml) or Isoniazid (1 µg/ml). Cells were washed with DMEM containing gentamycin (50 μg/ml) to remove extracellular bacteria and incubated for 24 hours. For CFU assay, macrophages were lysed using 0.1% Triton X-100 and serial dilutions of lysates were plated on 7H11 plates supplemented with 10% OADC (HIMEDIA, India) and 25 μg/ml kanamycin, 50 μg/ml hygromycin. Plates were incubated at 37°C and colonies were counted after 3-4 weeks as described earlier (Bisht et al. 2023).

### Flow cytometry

Approximately 0.5 – 1.0 million peritoneal macrophages from C57BL/6 mice were infected with various mycobacterial strains and either treated with 66.66 µg/ml of DQ-OVA (ThermoFisher Scientific, USA) for 60 minutes or stained with 500 ng/ml of Lysotracker Red (ThermoFisher Scientific, USA) and incubated for 45 minutes. Uninfected macrophages were served as a control group. The cells were scrapped, washed with FACS buffer and fluorescence was measured by flow cytometer.

### RNA sequencing analysis

RNA sequencing (RNA-seq) was carried out to analyze the transcriptomic changes in peritoneal macrophages from C57BL/6 mice in response to infections with various Mtb strains. In brief thioglycolate elicited peritoneal macrophages were either left uninfected or infected with wild-type H37Rv or H37Rv:*pe11*KO for 4 hours followed by washing to remove extracellular bacteria and then incubated for another 12 hours. Macrophages were harvested and total RNA was isolated using the Qiagen RNA extraction protocol and treated with DNase I to eliminate DNA contamination. The quality and integrity of the RNA samples were assessed using an Agilent Bioanalyzer, ensuring only high-quality RNA (RIN value of >7.0) was used for sequencing. Paired-end RNA sequencing was performed using an Illumina MiSeq Next Generation Sequencing system as per manufacturer’s instruction. The raw sequencing reads were filtered using the NGS QC toolkit to retain reads with a Phred score >33 and a read length of 150 bp. Additional quality checks and trimming of low-quality bases were performed using FASTQC and Trimmomatic, respectively, and the filtered reads were mapped to the reference genome of *Mus musculus*.

Differential gene expression analysis was performed using the Cufflinks tool, identifying genes with log2 fold changes ≤-1.00 as downregulated and ≥1.00 as upregulated. Functional analysis of differentially expressed genes (DEGs) was conducted using DAVID to annotate pathways and biological processes. Visualization of the results included enrichment analyses and heatmaps created using ggplot, while Venn diagrams were generated with ggVenndiagram to highlight overlapping gene sets.

### RNA isolation and real-time PCR (RT-PCR)

Total RNA was extracted from C57BL/6 peritoneal macrophages infected with various mycobacterial strains as well as from lung tissues harvested from C57BL/6 mice infected with various mycobacterial strains using Trizol reagent (Sigma-Aldrich, USA). RNA purity and concentration was assessed using a NanoDrop spectrophotometer (ThermoFisher Scientific, USA). The cDNA synthesis was carried out using Invitrogen SuperScript^TM^ III reverse transcriptase as per the manufacturer’s protocol. Real[time PCR was carried out using QuantStudio 5 Real[time PCR system (ThermoFisher Scientific, USA) with TB Green premix Taq (TAKARA) for detection. Expression of various genes was quantified by real-time PCR using β-actin as a reference gene. The primers used for RT-PCR are shown in Supplementary Table 2.

### Lactate estimation

For estimation of lactate, polar metabolite extractions from infected peritoneal macrophages from C57BL/6 mice were done using methods as previously described with minor modifications (Walvekar et al., 2018; Rajendran et al., 2022). Briefly, cell pellets were resuspended in 600 µl of 75% (v/v) ethanol with respective internal standard [2 nmol of D4-succinic acid]. The cell suspension was incubated at 80°C for 3 minutes with constant shaking and immediately kept on ice for 5 minutes. The mixture was then centrifuged at 20,000*g* for 10 minutes at 4°C, following which, the supernatant containing the desired polar metabolites was transferred to a new tube, dried under vacuum. For derivatization of the TCA intermediate metabolites, briefly, the dried extract was resuspended in 150 µl of water and they were extracted as reported previously (Walvekar et al., 2018; Rajendran et al., 2022) and stored at -80°C until LC-MS/MS analysis.

LC-MS runs were carried out using the AutoMS/MS acquisition method on an Agilent 6545 Q-TOF mass spectrometer fitted with an Agilent 1290 Infinity II UHPLC system as previously described with minor technical modifications (Walvekar et al., 2018; Rajendran et al., 2022). Reverse-phase LC separation was achieved using a Phenomenex Synergi Fusion-RP Column (150 mm X 4.6 mm, 4 μm, 80 Å) fitted with a Phenomenex guard column (3.2 mm X 8.0 mm). The solvents used for LC separation were buffer A: H_2_O + 0.1% HCOOH and buffer B: 50:50 H_2_O/MeOH + 0.1% HCOOH. LC-MS runs were 30 minutes post injection (for Derivatized Metabolites) as described previously (Walvekar et al., 2018; Rajendran et al., 2022). The temperatures of the autosampler (samples) and column were maintained 10 °C and 40 °C respectively. MS acquisition was performed using a dual Agilent jet stream electrospray ionization (AJS ESI) source in positive ionization mode, with the following MS parameters: Drying gas and sheath gas temperature at 320ִ°C, drying gas and sheath gas flow at 10 L/min, nebulizer pressure at 45 psi, capillary voltage (VCap) at 4000V, nozzle voltage at 1000V, and fragmentor voltage at 150V. Precursor selections were carried out with a maximum of 5 precursor ions per cycle with fixed collision energies set at 5 eV and 15 eV.

For data analysis, the data files were processed in Agilent MassHunter Qualitative Analysis 10.0, and all the peaks were manually validated based on relative retention times and fragments obtained, if any. All detected species were within a mass accuracy of 15 ppm and quantified by measuring the area under the curve (AUC) for different metabolites which were then normalized to levels of internal standard of the sample.

### Statistical analysis

For multiple group comparisons, one-way ANOVA test was performed. Calculations were performed using GraphPad Prism, version 9.1.2. Individual statistics of unpaired samples were performed using Student’s *t*-test. p < 0.05 was considered to be significant.

## Results

### PE11 regulates cell morphology and confers acid resistance to *M. tuberculosis*

We previously demonstrated that expression of Mtb PE11 in a surrogate bacterium *M. smegmatis* can alter its cell wall architecture (Singh et al., 2016). To further understand its role in the context of Mtb cell wall, we examined the morphology of *pe11* deleted *M. tuberculosis* strain (H37Rv:*pe11*KO) using scanning electron microscopy. It was found that the mutant strain displayed pronounced morphological aberrations, characterized by a significant reduction in cell size as compared to wild-type H37Rv and this morphological aberration can be rescued by complementing the mutant strain with a functional copy of the *pe11* gene (H37Rv:*pe11*KO::*pe11*) (Figures 1A–C). The morphological changes in the absence of *pe11* indicate that PE11 plays a critical role in maintaining the normal shape and size of Mtb. Since bacterial shape and size are known to influence survival in diverse environments as well as stress responses, these alterations may compromise its pathogenicity (Yang et al., 2016; Payros et al., 2021; Vijay et al. 2017).

**Figure 1.**
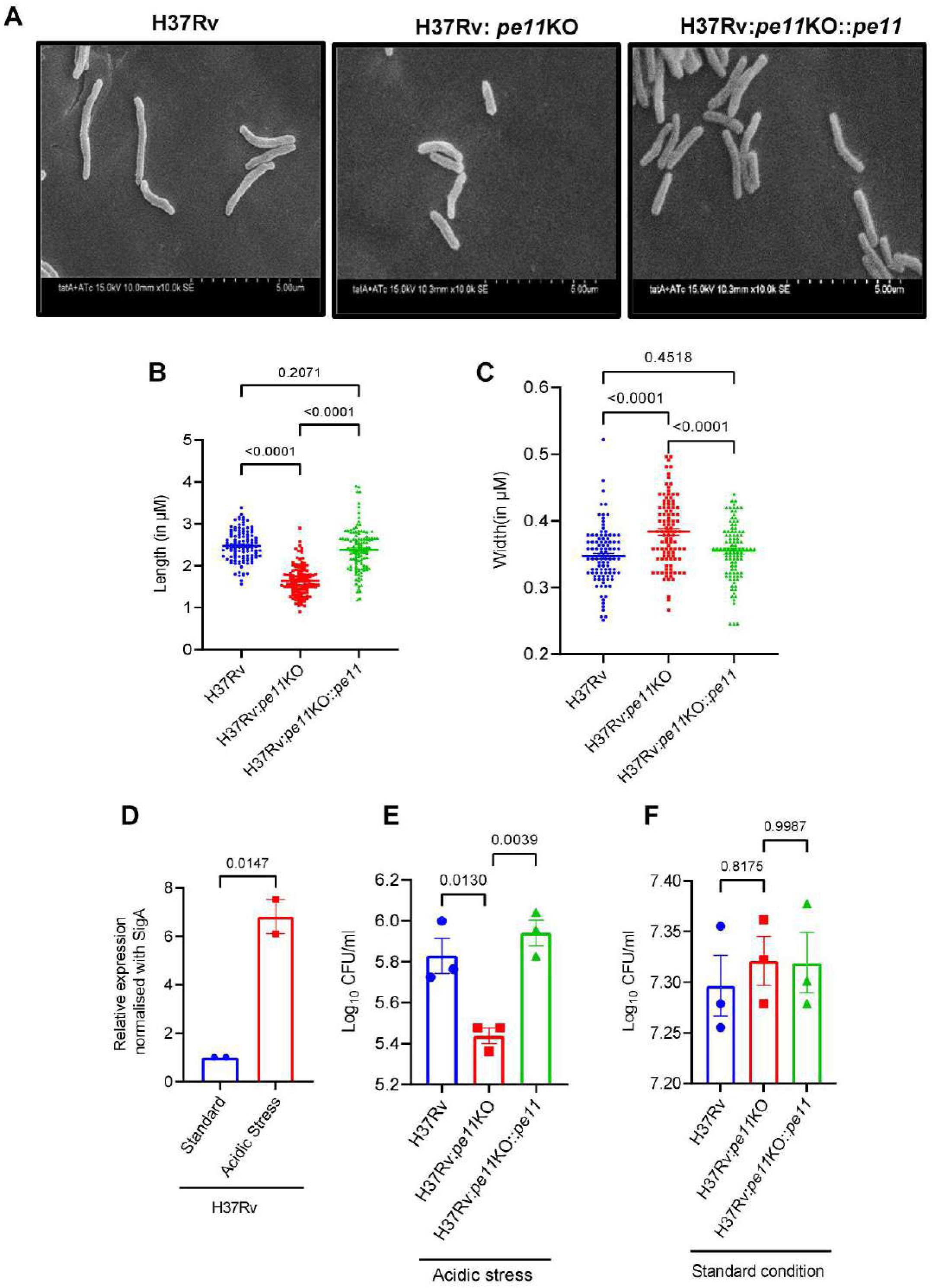
PE11 regulates mycobacterial cell morphology and confers resistance to acid stress. **(A)** Scanning electron microscopy (SEM) images of *M. tuberculosis* H37Rv, H37Rv:*pe11*KO, and complemented strain (H37Rv:*pe11*KO::*pe11*). Scale bar, 5 μm. Measurement of bacterial **(B)** length and **(C)** width from SEM images (n = 100 bacilli per strain). Data presented are mean ± SEM. **(D)** *M. tuberculosis* (H37Rv) bacteria were cultured in standard 7H9 medium or subjected to acidic (pH 4.5) stress, after which qPCR was performed to measure the transcript levels of *pe11. SigA* transcript level was used as an internal control. (**E**-**F**) Survival of different *M. tuberculosis* strains *viz* wild-type H37Rv, *pe11* knock-out H37Rv (H37Rv:*pe11*KO) and *pe11*-complemented *pe11*KO strain (H37Rv:*pe11*KO::*pe11*) under **(E)** Acidic stress (pH 4.5, 7 days), and **(F)** Standard culture conditions (7H9 medium, 3 days; control) was monitored. Data represent mean ± SEM of three independent experiments. Statistical analysis was performed using one-way ANOVA.

Expression of *pe11* was found to be upregulated during acidic stress, starvation and hypoxic conditions (Fisher et al., 2002; Voskuil et al., 2004; Schnappinger et al., 2003; Betts et al., 2002). Using *pe11*-null mutant of Mtb we further aimed to validate the expression of *pe11* under various stressors as well as evaluated its role in the bacterial survival. Interestingly, we found that the mRNA transcript level of *pe11* was significantly upregulated specifically in response to acidic stress condition as compared to standard culture condition in 7H9 medium (Figure 1D). It appears that PE11 significantly contributes to the bacterial adaptation to acidic stress conditions, since *pe11*-null mutants survived poorly under this condition as compared to the wild-type strain. Interestingly, the *pe11*-null mutants could regain their ability to survive when these mutants are complemented with a functional copy of *pe11* gene (Figure 1E), However no significant differences in bacterial survival was observed among the wild-type H37Rv, H37Rv:*pe11*KO and H37Rv:*pe11*KO::*pe11* strains when grown in standard 7H9 medium (Figure 1F). This suggests that PE11 plays a critical role in the survival of Mtb under stressful environments, which the bacilli encounter during infection inside the phagosome of host macrophages (Rai et al., 2022).

### PE11 exacerbates inflammation during mycobacterial infection and provides survival advantage to the bacilli *in vivo*

Earlier, Mtb PE11 was found to confer higher resistance to acidic stress, antibiotics and survival advantage to recombinant *M. smegmatis* expressing this protein (Singh et al., 2016; Dahiya et al., 2024). In addition, *M. smegmatis* expressing PE11 was found to induce production of pro-inflammatory cytokines in macrophages (Singh et al., 2016, Deng et al., 2015). To further validate these observations, we examined the inflammatory properties of PE11 *in vivo* using a virulent *M. tuberculosis* strain H37Rv and corresponding *pe11*-null mutant (H37Rv:*pe11*KO). Accordingly, C57BL/6 mice were infected *via* aerosol route with low-dose of H37Rv, H37Rv:*pe11*KO and H37Rv:*pe11*KO::*pe11* (which is regarded as a more realistic animal model of human tuberculosis, Plumlee et al., 2021). Expectedly, mice infected with H37Rv:*pe11*KO showed significantly reduced lung bacillary loads compared to wild-type and complemented strains at 30 days post-infection (dpi) (Figure 2A). Mice infected with wild-type H37Rv and complemented H37Rv:*pe11*KO::*pe11* developed well-organized granulomas by 30 days, while H37Rv:*pe11*KO-infected lungs exhibited sparse, disorganized lesions as suggested by haematoxylin and eosin staining of mice lung sections (Figure 2B). To further understand PE11’s role in driving inflammation, we assessed the serum levels of key pro-inflammatory cytokines, mainly, IL-1β, IL-6 and TNF-α in these mice. Interestingly, mice infected with H37Rv and H37Rv:*pe11*KO::*pe11* exhibited significantly higher levels of these pro-inflammatory-type cytokines compared to those infected with H37Rv:*pe11*KO at 30 dpi (Figure 2C-E). These cytokines were also found to be higher in lung tissues of mice infected with H37Rv and H37Rv:*pe11*KO::*pe11* as compared to mice infected with H37Rv:*pe11*KO (Figure 2F-H). Given that macrophages serve as a critical niche for Mtb survival, providing both a replicative environment and a platform for immune modulation (Cohen et al., 2018), cytokine levels were further assessed following Mtb infection in macrophages. Consistent with the *in vivo* results, H37Rv- and H37Rv:*pe11*KO::*pe11*-infected macrophages showed significantly higher levels of IL-1β, IL-6 and TNF-α cytokines when compared with macrophages infected with H37Rv:*pe11*KO (Figure 2I-K).

**Figure 2.**
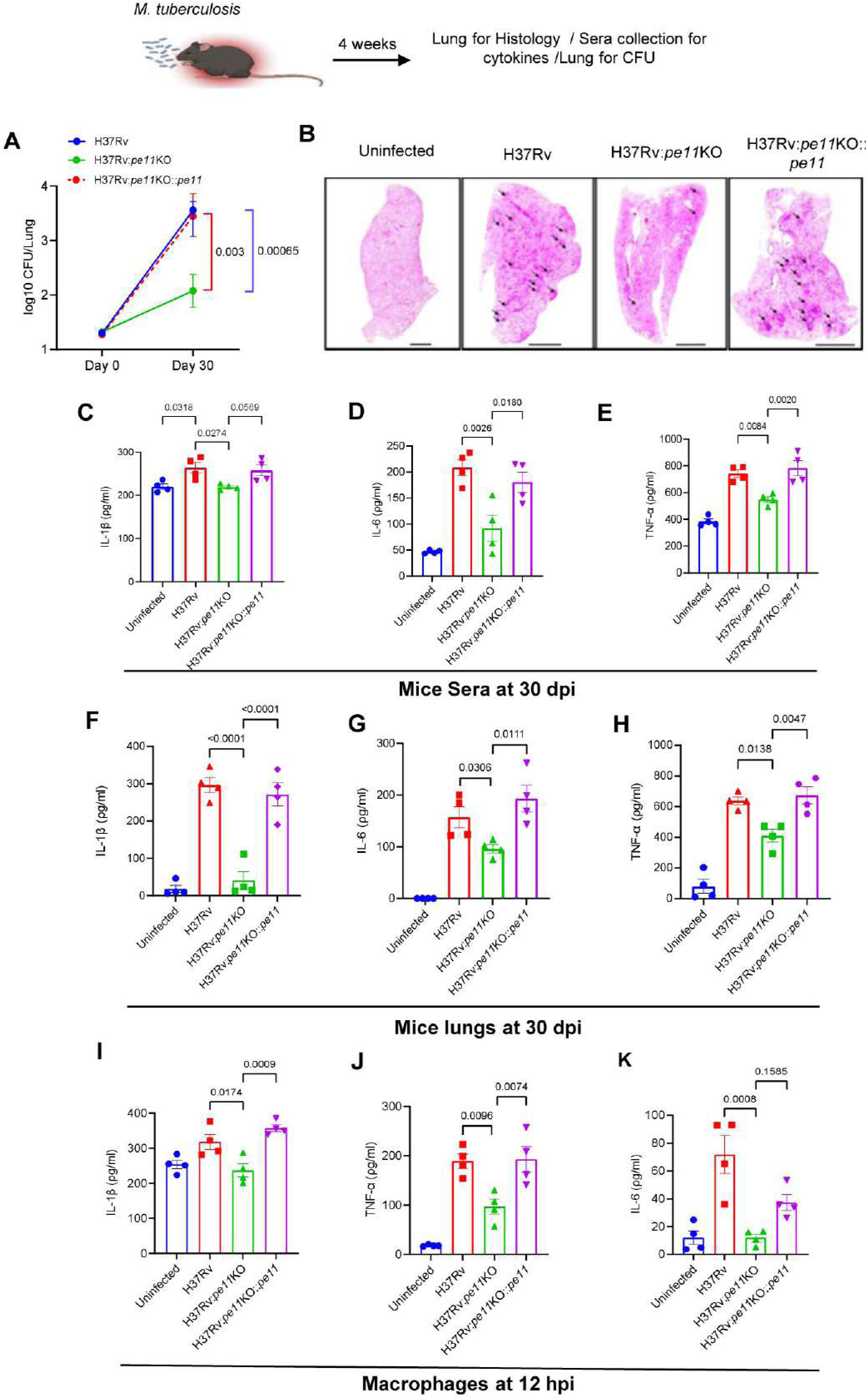
PE11 provides survival advantage and promotes inflammation during *M. tuberculosis* infection. **(A)** Bacterial burden in the lungs of C57BL/6 mice infected *via* aerosol with various *M. tuberculosis* strains (H37Rv, H37Rv:*pe11*KO, or complemented strain [H37Rv:*pe11*KO::*pe11*]) at 30 days post-infection (dpi). **(B)** Hematoxylin and Eosin**-**stained lung sections showing granuloma-like structures (arrows). Scale bar, 500 μm (overview, 4X) and 200 μm (10X). Levels of IL-1β, IL-6, TNF-α cytokines in **(C-E)** the sera and **(F-H)** lung lysates from mice infected with various strains of *M. tuberculosis* (Dara presented as mean ± SEM, n = 4). **(I-K)** Levels of IL-1β, IL-6 and TNF-α cytokines secreted by C57BL/6 peritoneal macrophages infected with various *M. tuberculosis* strains at 12 hours post-infection (hpi, MOI 5). Data presented as mean ± SEM (n = 4). Statistical significance was determined by one-way ANOVA. Uninfected mice were used as control in all the experiments.

To further confirm the inflammatory role of Mtb PE11, C57BL/6 mice were infected with 50 million CFU of *M. smegmatis* harboring either backbone control vector (*Msmeg*-pVV) or wild-type Mtb PE11 (*Msmeg*-PE11) and blood profiling was carried out at day 3 days post-infection. It was found that infection with *Msmeg*-PE11 resulted in a pronounced inflammatory response characterized by an increase of neutrophils and monocytes (Supplementary Figure 1A and B). In addition, exacerbated lung pathology and higher recruitment of CD45^+^ macrophages were observed in the lung of *Msmeg-*PE11 infected mice as compared to those infected with *Msmeg*-pVV (Supplementary Figure 1C, D).

These findings suggest that the PE11 protein plays an important role in granuloma formation and drives pro-inflammatory responses during infection, which is a key feature of TB pathogenesis (Cronan et al., 2022; McCaffrey et al., 2022; Rubin EJ, 2009; Etna et al., 2014; Elkington et al., 2022).

### PE11 modulates the transcriptomic landscape in primary macrophages during infection with Mtb

The above findings as well as by others (Singh et al., 2016) suggest that that Mtb PE11 plays a significant role in remodelling the mycobacterial cell wall and regulate the inflammatory milieu during Mtb infection to favor Mtb survival. Therefore, PE11 appears to be pleiotropic in nature as many other members of the PE/PPE family proteins (Ahmed et al., 2015; Akhter et al., 2012; Mukhopadhyay and Balaji, 2011). Therefore, we speculated that PE11 may possess multiple functions to regulate many other host functions. To further investigate how PE11 influences overall host immune functions to favour its survival, we performed transcriptome analysis in the presence or absence of PE11 during infection of peritoneal macrophages isolated from C57BL/6 mice with H37Rv or H37Rv:*pe11*KO for 12 hours. It was found that, PE11 causes widespread transcriptional changes to significantly reprograms the macrophage immune responses (Figure 3). Infection with wild-type H37Rv resulted in significant transcriptional reprogramming, with significant upregulation of 2,167 genes and downregulation of 1,471 genes compared to uninfected controls (Figure 3A). In contrast, infection of macrophages with H37Rv:*pe11*KO strain resulted in upregulation of 894 genes and downregulation of 282 genes relative to the uninfected macrophages. Direct comparison between H37Rv:*pe11*KO and H37Rv-infected macrophage transcriptome showed upregulation of 471 genes and downregulation of 299 genes (Figure 3A), emphasizing PE11’s important role in shaping the macrophage transcriptional responses upon infection. Significantly differentially expressed genes (DEGs) were identified using a stringently adjusted p value (padj ≤ 0.05) and log2 fold-change cutoff value of greater than ±1.0 (Figure 3B). These DEGs between H37Rv- and H37Rv:*pe11*KO-infected macrophages were further used for Reactome enrichment and KEGG pathway analyses (Figure 3C, D). Heatmap of the top 50 DEGs are shown in Supplementary Figure 2.

**Figure 3.**
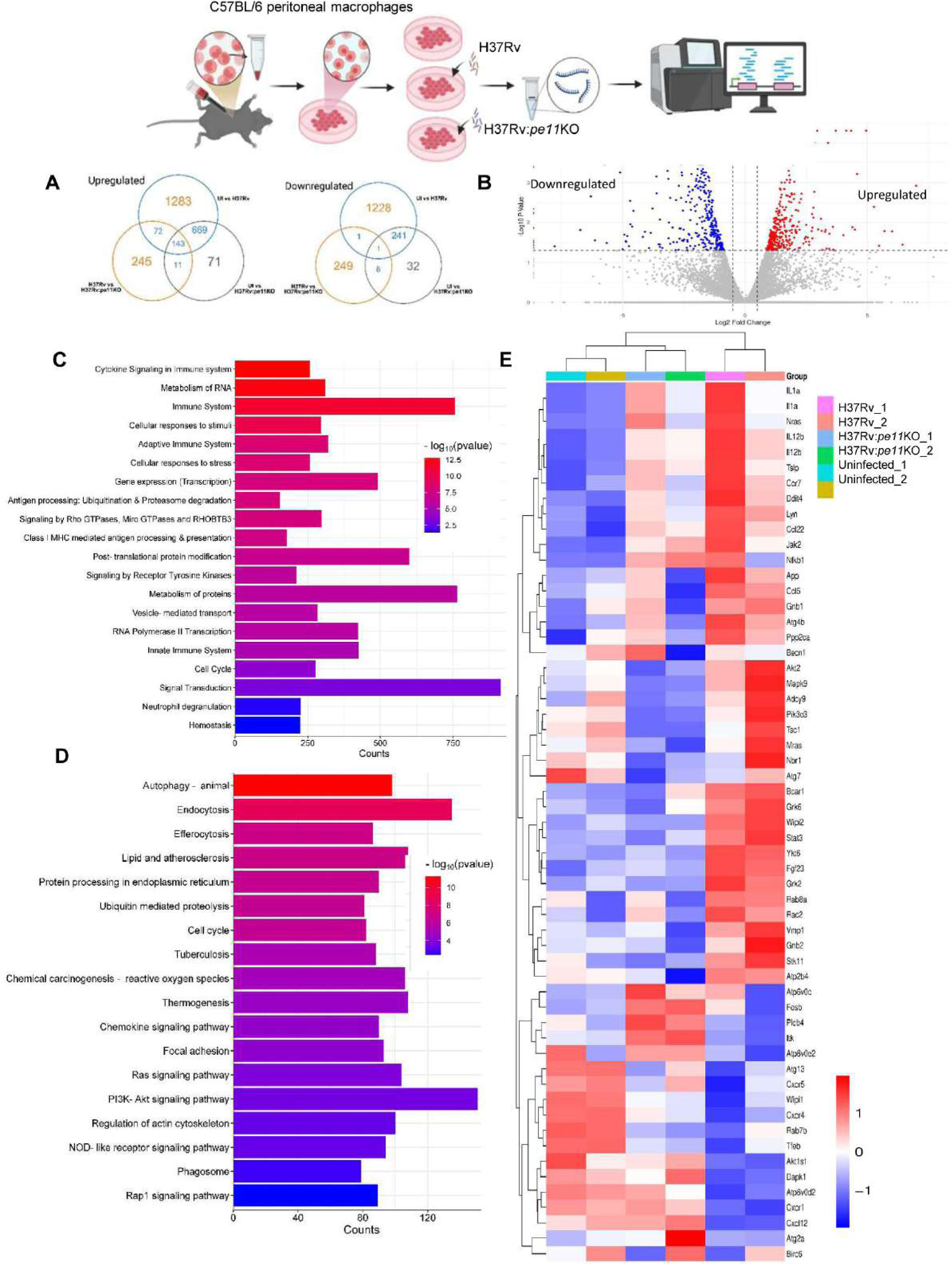
PE11 reprograms the host transcriptional responses to infection. **(A)** Venn diagram of differentially expressed genes (DEGs) in C57BL/6 peritoneal macrophages at 12 hours post-infection with H37Rv, H37Rv:*pe11*KO or uninfected control (UI) (*padj* < 0.05, log2 Fold Change (FC) > 1 or < 1). **(B)** Volcano plot of DEGs in H37Rv-versus H37Rv-*pe11*KO-infected macrophages. Red: upregulated (log2FC > 1); blue: downregulated (log2FC < −1); gray: nonsignificant. **(C)** Reactome pathway enrichment of DEGs from H37Rv versus H37Rv-*pe11*KO. Top pathways include immune signaling and metabolic processes. **(D)** KEGG pathway analysis of DEGs highlighting autophagy, lysosomal function, and inflammatory responses. **(E)** Heatmap of DEGs (rows) across infected macrophages (columns), clustered by expression patterns in immune response, autophagy, and lysosomal pathways. Color scale: log2-normalized counts.

Gene ontology enrichment analyses indicate that the differentially expressed genes were predominantly associated with immune pathways critical for bacterial clearance, including those involved in cytokine signalling, and innate and adaptive immune responses (Figures 3C). KEGG pathways analysis further revealed that the genes associated with autophagy, endocytosis and lipid metabolism were differentially regulated between wild-type H37Rv- and H37Rv:*pe11*KO-infected macrophages (Figure 3D), indicating that PE11 influences both the immune and metabolic functions during macrophage infection. Autophagy is known to prevent excessive inflammation and acts as a cell-autonomous defence against intracellular pathogens including Mtb (Yuk et al., 2012; Deretic, 2014; Kim et al., 2019; Kimmey and Stallings, 2016), and its connection with endocytosis pathway plays a critical role in clearing engulfed bacteria through phagosomal maturation (Gray et al., 2016; Birgisdottir and Johansen, 2020). Since both these pathways utilize lysosomes as common endpoint to clear bacterial infection (Birgisdottir and Johansen, 2020), we speculated that PE11 may be involved in inhibition of autophagy by regulating genes associated with lysosome biogenesis and acidification. Analyses of the genes involved in autophagy like ATG2a, ATG7, ATG13, BECN1, STK11), endocytosis (GRK2, GRK6, HRAS, RAB8A, TRAF6) and lysosomal biogenesis and acidification (TFEB, Atp6v0c, Atp6v0e2), revealed distinct expression patterns in the heatmap (Figure 3E). Expression of autophagy associated genes like ATG2a, ATG13, were found to be muted in wild-type Mtb infected macrophages compared to those infected with *pe11*KO strains. Interestingly, expression of TFEB, a key transcription factor associated in regulation of autophagy and lysosome biogenesis (Napolitano and Ballabio, 2016), was found to be downregulated in macrophages infected with wild-type Mtb compared to the *pe11*-null mutant as well as uninfected macrophages (Figure 3E). Pro-inflammatory cytokines like IL-6 and IL-1β were significantly downregulated in *pe11*KO strain, while chemokine receptors (Cxcr1, Cxcr4) were partially upregulated (Figure 3E). The transcription factor Nfkb1 was upregulated in wild-type Mtb infections, while Stat3 and Jak2 were downregulated in *pe11*KO, suggesting that PE11 may modulate pro-inflammatory cytokine signalling (Figure 2 and 3C-E). It appears that deletion of *pe11* resulted in a shift towards enhanced autophagy that potentially aids in bacterial clearance and reduced inflammation indicating that PE11 probably plays vital roles in blocking the host immune defence mechanisms like autophagy and promotes inflammatory responses.

### PE11 suppresses autophagy and antigenic-degradation during infection

Our previous studies demonstrated that Mtb PE11 protein confers a survival advantage to the mycobacteria (Singh et al., 2016; Dahiya et al., 2024, Figure 2A). Transcriptome analysis identified several genes related to autophagy and lysosomal biogenesis were altered by PE11, suggesting a potential role for PE11 in manipulating host autophagy process and lysosomal biogenesis and acidification. Mtb has evolved mechanisms to subvert host immune defence mechanisms including autophagy, a critical pathway for elimination of intracellular pathogens (Chandra et al., 2022; Castillo et al., 2012; Kimmey et al., 2015; Typas, 2023). By disrupting autophagosome formation and lysosomal maturation and autophagic degradation, Mtb ensures its survival and replication within the hostile intracellular environment (Golovkine et al., 2023, Deretic, 2014).

In the previous section, PE11 was found to alter the expression of key autophagy associated genes like *atg5*, *becn1* (Yang et al., 2021). Given the central role of ATG5 and Beclin1 (encoded by BECN1 gene) in autophagosome formation (Changotra et al., 2022; Bradfute et al., 2013; Kang et al., 2011), we speculated that PE11 targets the autophagic pathway to evade the host immunity. Accordingly, the transcript levels of these genes were validated in macrophages infected with wild-type Mtb and *pe11*-complemented *pe11*KO strains (*pe11*KO::*pe11*) and compared their levels in macrophages infected with *pe11*-deficient *pe11KO* strain (Figure 4A, B). It was found that, macrophages infected with *pe11*KO strain had significantly higher levels of *becn1* and *atg5* transcripts compared to those of macrophages infected with wild-type Mtb or *pe11-*complemented *pe11*KO strain. Similar results we obtained *in vivo*, with lungs from mice infected with wild-type and *pe11*-complemented strain showing decreased *atg5* and *becn1* expression compared to those infected with *pe11*KO strain (Figure 4C, D).

**Figure 4.**
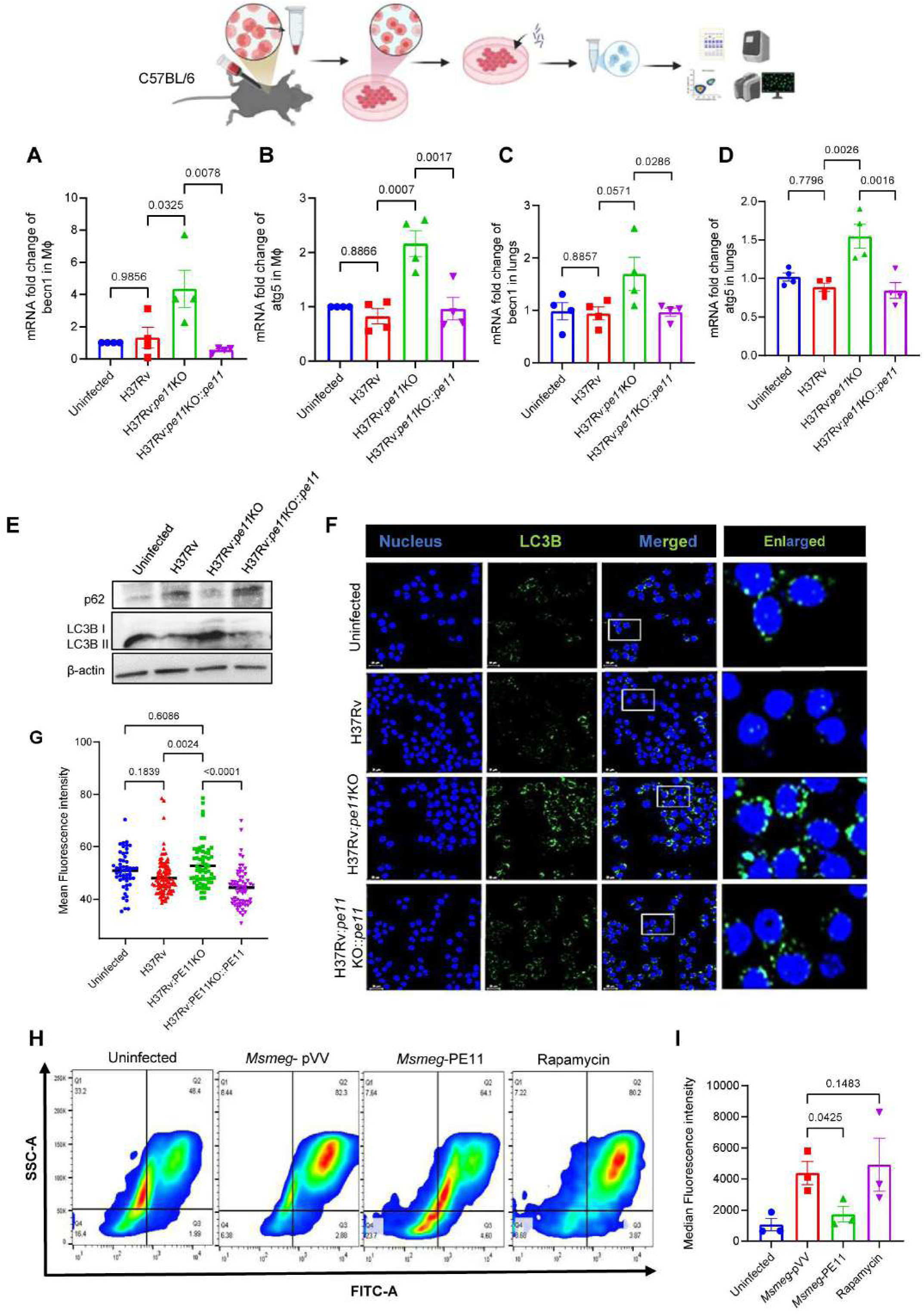
PE11 impairs autophagy and proteolytic degradation during *M. tuberculosis* infection. **(A, B)** Fold changes in the mRNA levels of *BECN1* (**A**) and *ATG5* (**B**) in *M. tuberculosis*-infected C57BL/6 peritoneal macrophages (MOI 5, 12 hours post-infection (hpi); mean ± SEM, n = 3 experiments). **(C, D)** Fold changes in the mRNA levels of *BECN1* (**C**) and *ATG5* (**D**) in lung tissue from C57BL/6 mice infected with low-dose *M. tuberculosis* strains (20 CFU, 30 days post-infection; mean ± SEM, n = 4 mice). **(E)** Measurement of LC3B and p62 protein levels by Western Blotting in *M. tuberculosis*-infected C57BL/6 macrophages. **(F)** Confocal microscopy of LC3B puncta (red) in BMC2 macrophages infected with various *M. tuberculosis* strains (DAPI: blue; scale bar: 10 μm) and (**G**) quantitation of LC3B+ puncta per cell (mean ± SEM, 100 cells/condition). **(H)** Measurement of fluorescence associated with proteolytic degradation of DQ-OVA using flowcytometry in macrophages infected with *Msmeg-*pVV and *Msmeg-*PE11 (MOI 1:10, 6 hpi). (**I**) Median fluorescence intensities of different experimental groups were calculated and presented as mean ± SEM of 3 independent experiments. Statistical significance was determined by one-way ANOVA. Uninfected cells were used as control in all the experiments.

Next, we assessed the extent of autophagic flux in macrophages infected with either *Msmeg*-PE11/*Msmeg*-pVV or H37Rv/H37Rv:*pe11*KO/H37Rv:*pe11*KO::*pe11* and measured the expression levels of autophagy markers, like p62 and LC3BII. The level of lipidated form of LC3B, also known as LC3BII, is commonly regarded as a key indicator of autophagosome formation, while accumulation of p62 reflects impaired autophagosome degradation (Mizushima et al., 2010). Interestingly, it could be observed that infection of macrophages with H37Rv/H37Rv:*pe11*KO::*pe11* resulted in a significant reduction of LC3BII levels and an accumulation of p62 as compared to macrophages infected with H37Rv:*pe11*KO (Figure 4E). Similar results were observed in macrophages infected with *Msmeg*-pVV when compared to those infected *Msmeg*-PE11 (Supplementary Figure 3A-C). Immunofluorescence analysis further confirmed this observation, as PE11-expressing strains showed diminished LC3B puncta formation (Figure 4F, G and Supplementary Figure 4D), indicating suppression of autophagic initiation. Since lysosomal degradation of antigen is a critical step in the autophagic process (Øynebråten et al., 2022; Kimmey and Stallings, 2016; Bradfute et al., 2013, Yang and Wang, 2021), we speculated that PE11 may also hinder lysosomal antigenic degradation. To examine this hypothesis, peritoneal macrophages isolated from C57BL/6 mice were infected with either Msmeg-pVV or Msmeg-PE11, and antigen degradation was assessed using DQ-OVA. DQ-OVA is a self-quenched, fluorescently labelled ovalbumin that emits green fluorescence upon proteolytic degradation. Using flow cytometry, when the fluorescence emitted was quantified, it was found that the macrophages infected with *Msmeg*-PE11 displayed significantly reduced DQ-OVA fluorescence compared to those infected with *Msmeg*-pVV control (Figure 4H, I), indicating that PE11 impairs lysosomal degradative function.

Taken together, these findings provide evidence that PE11 interferes with autophagic influx as evidenced by observed reduction of LC3BII levels and increased accumulation of p62, by inhibiting autophagy-associated gene expression and impairing antigenic degradation. These effects likely represent a hitherto unknown virulence mechanism of PE11 that promotes mycobacterial survival by evading host autophagic degradation.

### PE11 impairs lysosomal function by inhibiting its acidification

In the previous section, we discovered that PE11 impairs lysosomal degrative function which could be due to inefficient lysosomal acidification as lysosomal enzymes require acidic pH for their optimal proteolytic activities (Ballabio and Bonifacino, 2020; Lawrence and Zoncu, 2019). Therefore, we measured the lysosomal pH using LysoTracker Red, a pH-sensitive fluorescent dye. It was found that, macrophages infected with *Msmeg*-PE11 had significantly lower fluorescence intensity as compared to those infected with *Msmeg*-pVV (Figure 5A, 5B) indicating reduced acidification in lysosomes in the presence of PE11. A similar trend was observed in macrophages infected with wild-type Mtb H37Rv or *pe11*KO strain complemented with *pe11* (H37Rv:*pe11*KO::*pe11*), both of which showed lower fluorescence compared to the PE11-deficient H37Rv:*pe11*KO strain (Figure 5C, D). These findings suggest that PE11 impairs lysosomal acidification and thereby potentially compromising its proteolytic degradative function.

**Figure 5.**
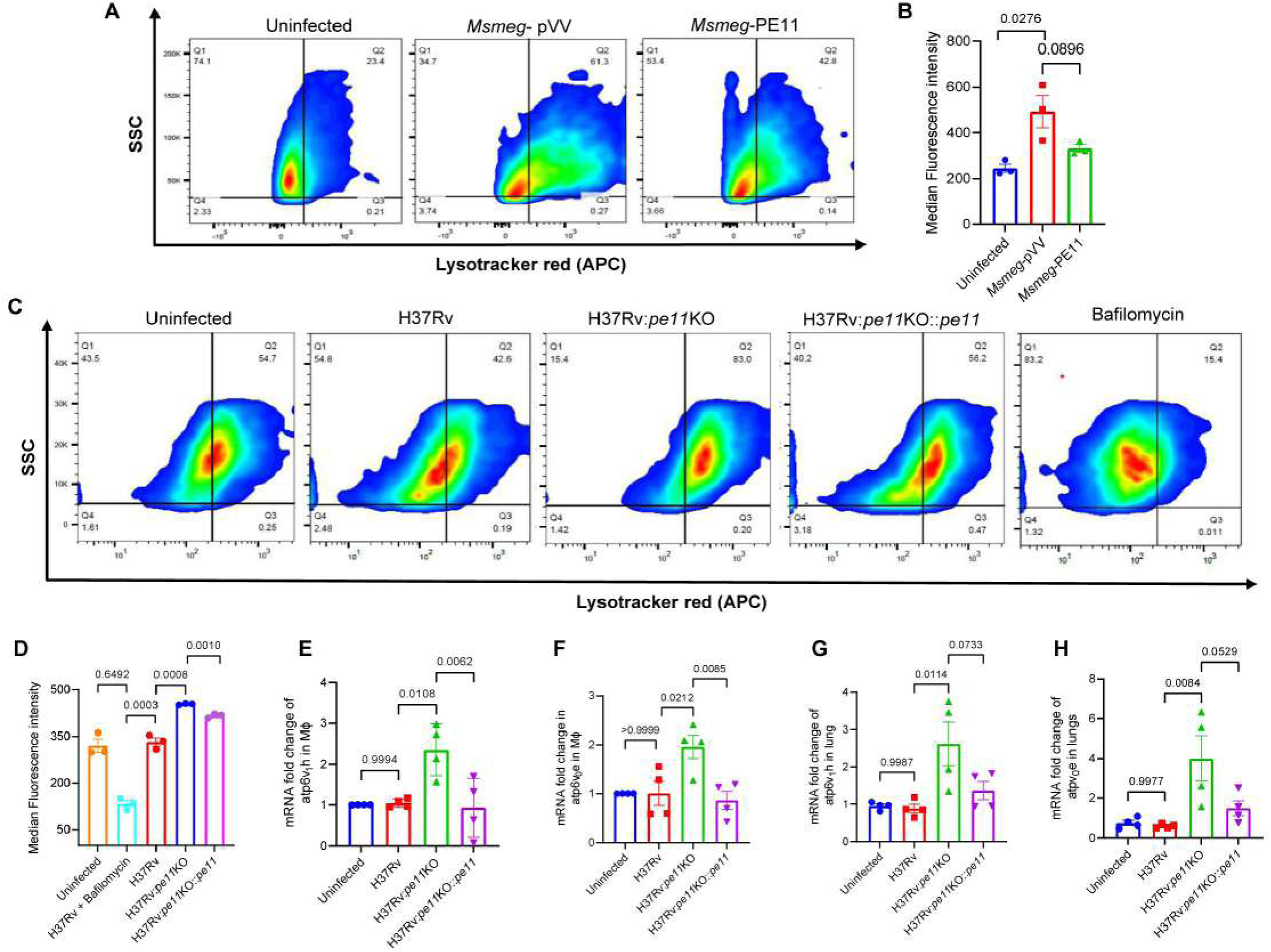
PE11 impairs lysosomal acidification by inhibiting expression of V-ATPase subunits. **(A, B)** Flow cytometric measurement of lysosomal acidification using LysoTracker Red in C57BL/6 peritoneal macrophages infected with either *Msmeg*-pVV or *Msmeg-*PE11 at MOI 1:10 for 4 hours. Median fluorescence intensities were calculated and presented as mean ± SEM of 3 independent experiments (C, D). Flow cytometric measurement of lysosomal acidification using LysoTracker Red in C57BL/6 peritoneal macrophages infected with various *M. tuberculosis* strains (H37Rv/H37Rv:*pe11*KO/H37Rv:*pe*11KO::*pe11*) infected at MOI 1:5 for 12 hours. Median fluorescence intensities of different experimental groups were calculated and presented as mean ± SEM (n = 3). **(E, F)** Fold changes of mRNA levels of *ATPV1h* and *ATPV0E* genes in *M. tuberculosis*-infected C57BL/6 peritoneal macrophages at 12 hours post-infection (MOI 5, 12 hours post-infection (hpi)). **(G, H)** Fold changes of mRNA levels of *ATPV1h* and *ATPV0E* genes in the lung tissue obtained from mice infected with low-dose *M. tuberculosis* strains (20 CFU, 30 days post-infection). Data represent mean ± SEM (n = 3 experiments for **E** and **F**; n = 4 mice for **G** and **H**). Statistical significance was determined by one-way ANOVA

To further elucidate the mechanism behind impaired lysosomal acidification, we examined the expression of Vacuolar-type ATPase (V-ATPase) subunits, proton pumps critical for lysosomal acidification (Mindell, 2012; Bouché et al., 2016; Colacurcio and Nixon, 2016; Song et al., 2020; Chen et al., 2025). V-ATPases acidify lysosomes by translocating protons across the lysosomal membrane, a process that consumes ATP (Perera and Zoncu, 2016). Expectedly, the transcripts of two essential subunits, Atp6v1h (Yambire et al., 2019) and Atp6v0e (Sun et al., 2020), were found to be significantly downregulated in macrophages infected with wild-type Mtb or the *pe11*-complemented *pe11*KO strains compared to those infected with the *pe11*KO strain (Figure 5E, F) as also observed in our transcriptome data (Figure 3E). This downregulation was also evident *in vivo*, as lungs from wild-type Mtb-infected mice exhibited significantly reduced expression of *Atp6v1h* and *Atp6v0e* when compared with those infected with the *pe11*-deficient Mtb (Figure 5G, H). These findings suggest that PE11 interferes with lysosomal acidification by suppressing V-ATPase subunit gene expression. Therefore, it appears that PE11 contributes to the pathogen’s ability to evade host defence mechanisms by impairing cellular degradative processes (Saftig and Klumperman et al., 2009; Perera and Zoncu, 2016).

### PE11 exploits the FLCN-lactate-TFEB signalling axis during infection

Interestingly, comparison of the transcriptomic data between macrophages infected with wild-type H37Rv and those infected with *pe11*-deficient strain (H37RV:*pe11*KO) revealed that, PE11 also suppressed the expression of Transcription Factor EB (TFEB) mRNA levels (Figure 3E). As TFEB is a key regulator of many genes involved in the autophagy and lysosomal biogenesis pathway, including *atg5*, *becn1*, and *V-ATPase* subunit genes (Sardiello et al., 2009; Settembre et al., 2011; Cortes et al., 2019; Song et al., 2021; Yang and Wang, 2021; Gros and Muller, 2023; Medina et al., 2011; Settembre et al., 2012), its downregulation provides a plausible explanation for the impaired autophagy and lysosomal acidification observed during infection with PE11-expressing strains. When we validated the transcriptome data by qRT-PCR, it was found that TFEB mRNA levels were significantly reduced in macrophages infected with H37Rv compared to those infected with *pe11*-knockout strain (H37Rv*-pe11KO*) (Figure 6A). This was consistent with the observations in lung tissues of infected mice, where TFEB mRNA levels were also significantly lower in H37Rv-/H37Rv-*pe11*KO::*pe11*-infected mice compared to H37Rv-*pe11*KO-infected mice, further corroborated a role of PE11 in suppressing TFEB activity during infection (Figure 6B). TFEB activity was further assessed using confocal microscopy, which showed that PE11 reduced overall expression as well as nuclear translocation of TFEB. Macrophages infected with PE11-expressing strains (H37Rv or H37Rv:*pe11*KO::*pe11*) showed significantly reduced total and nuclear TFEB levels as compared to those infected with the PE11-deficient strain (H37Rv:*pe11*KO) (Figure 6C–E).

**Figure 6.**
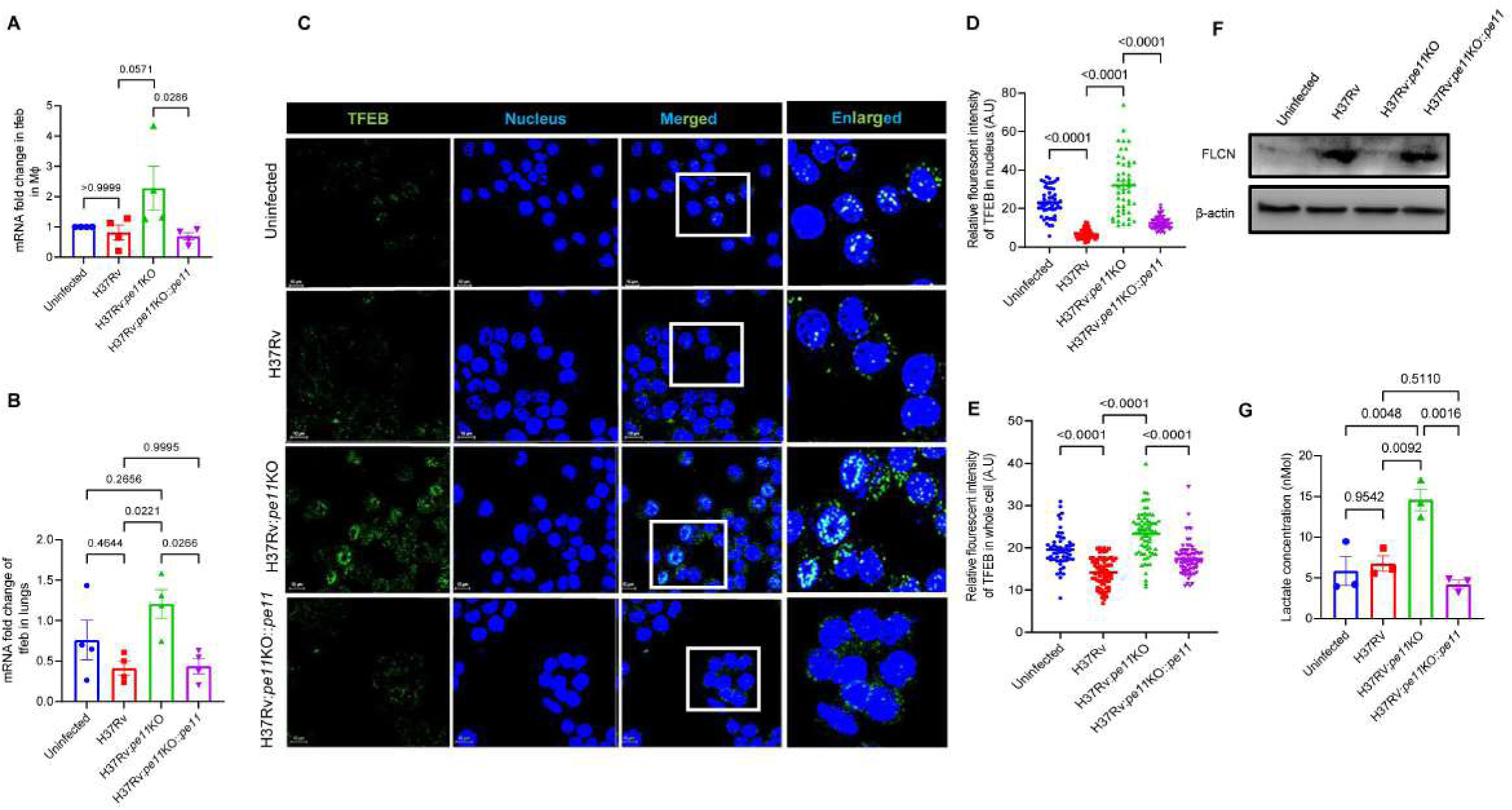
PE11 modulates FLCN-lactate-TFEB axis during *M. tuberculosis* infection. **(A, B)** Fold changes of TFEB mRNA levels in (**A**) peritoneal macrophages infected with various strains of *M. tuberculosis* (MOI 5) at 12 hours post-infection (hpi)) and (**B**) in lung tissues from mice either left uninfected or infected with various strains of *M. tuberculosis* strains (20 CFU) at 30 days post-infection (dpi). **(C, E)** Subcellular localization of TFEB in BMC2 macrophages infected with various strains of *M. tuberculosis* at MOI 1:10. (**C**) Representative immunofluorescence images (TFEB: green; DAPI: blue; scale bar: 10 μm), with quantification of (**D**) total cellular and (**E**) nuclear TFEB intensity (n = 50 cells). **(F)** FLCN protein levels measured by Western blotting with β-actin as loading control in C57BL/6 macrophages infected with various *M. tuberculosis* strains. **(G)** Lactate levels as measured by LC/MS in C57BL/6 macrophages infected with various *M. tuberculosis* strains at 12 hpi. Data represent mean ± SEM. Statistical significance was determined by one-way ANOVA. Cells left uninfected were used as control in all the experiments.

It has been shown that nuclear accumulation and activation of TFEB transcription factor is regulated by cellular Folliculin (FLCN) level (Paquette et al., 2021). By acting as a negative regulator upstream of TFEB, FLCN plays a crucial role in controlling lysosome function and host immune responses (El-Houjeiri et al., 2019, Napolitano et al., 2022). Therefore, we investigated whether PE11 modulates FLCN to inhibit TFEB activity. Interestingly, FLCN protein level was found to be significantly upregulated in macrophages infected with H37Rv or H37Rv:*pe11*KO::*pe11* strains compared to those infected with *pe11*-deficient H37Rv:*pe11*KO Mtb strain (Figure 6F). Interestingly, FLCN is also known to be an uncompetitive inhibitor of lactate dehydrogenase A (LDHA), a glycolytic enzyme, in addition to its canonical function to regulate mTOR (Woodford et al., 2021). In another interesting study, it has been reported that lactate can stabilize TFEB by inhibiting its proteolytic degradation through lactylation of lysine residues (Huang et al., 2024). Therefore, we compared the cellular lactate levels in macrophages infected with H37Rv, H37Rv:*pe11*KO::*pe11* and H37Rv:*pe11*KO, and found that cellular lactate levels are significantly reduced in macrophages infected with PE11-expressing strains (H37Rv and H37Rv:*pe11*KO::*pe11*) compared to H37Rv:*pe11*KO (Figure 6G). Thus, these findings suggest that PE11 employs a dual mechanism to impede host autophagy. Firstly, it upregulates FLCN to suppress TFEB activity and secondly, through FLCN-mediated inhibition of LDHA, it reduces lactate levels to prevent TFEB lactylation and stabilization. This coordinated disruption of lysosomal function and autophagic flux, highlight PE11 as a critical factor that aids Mtb to evade host immune mechanisms. By manipulating the host’s FLCN-lactate network, PE11 exemplifies how pathogens can rewire host cellular machinery for its survival.

### Lactate supplementation rescues lysosomal function through TFEB stabilization and promotes Mtb clearance

The data presented in the previous section suggest that PE11-mediated subversion of autophagy is associated with decreased lactate production that affects TFEB stability. Therefore, we speculated that, exogenous lactate supplementation may restore TFEB stability and thereby lysosome function to promote mycobacterial clearance from infected cells. Therefore, macrophages were infected with H37Rv in the absence or presence of 10 mM lactate, at a concentration found to be non-cytotoxic (Supplementary Figure 4A, B). Interestingly, it was observed that, treatment of macrophages with 10 mM lactic acid could significantly reduce mycobacterial burdens at 24 hours and 48 hours post-infection (Figure 7A). However, lactic acid at 10 mM and 20 mM had no direct bactericidal effect in Mtb axenic culture (Figure 7B) indicating that the anti-mycobacterial effect lactate is due to modulation of host cell functions essential for Mtb survival inside macrophages. Lactate treatment was found to be associated with increased accumulation of TFEB (Figure 7C, D) which could be due to increased stability of TFEB due to increased lactylation by the supplemented lactate (Huang et al., 2024). Lactate supplementation was also found to be associated with improved lysosomal acidification in Mtb-infected macrophages treated with lactate as compared to Mtb-infected macrophages left untreated (Figure 7E). When C57BL/6 mice were given a single dose of 500 mg/kg lactate at day 2 post infection with *Msmeg*-pVV/*Msmeg*-PE11, it was found that, mice infected with *Msmeg*-PE11 had higher bacterial load in lungs as compared to *Msmeg*-pVV (Supplementary Figure 5) which was expected (Singh et al., 2016). However, lactate supplementation decreased bacterial load in mice infected with *Msmeg*-PE11 as well as *Msmeg*-pVV (Supplementary Figure 5). Importantly, when lactate supplementation was used in combination with frontline anti-TB drugs. like Isoniazid and Rifampicin, it was found that lactate significantly improved the efficacy of Isoniazid/Rifampicin in clearing Mtb bacilli (Figure 6E), supporting the potential of lactate as an adjunctive therapy against Mtb infection.

**Figure 7.**
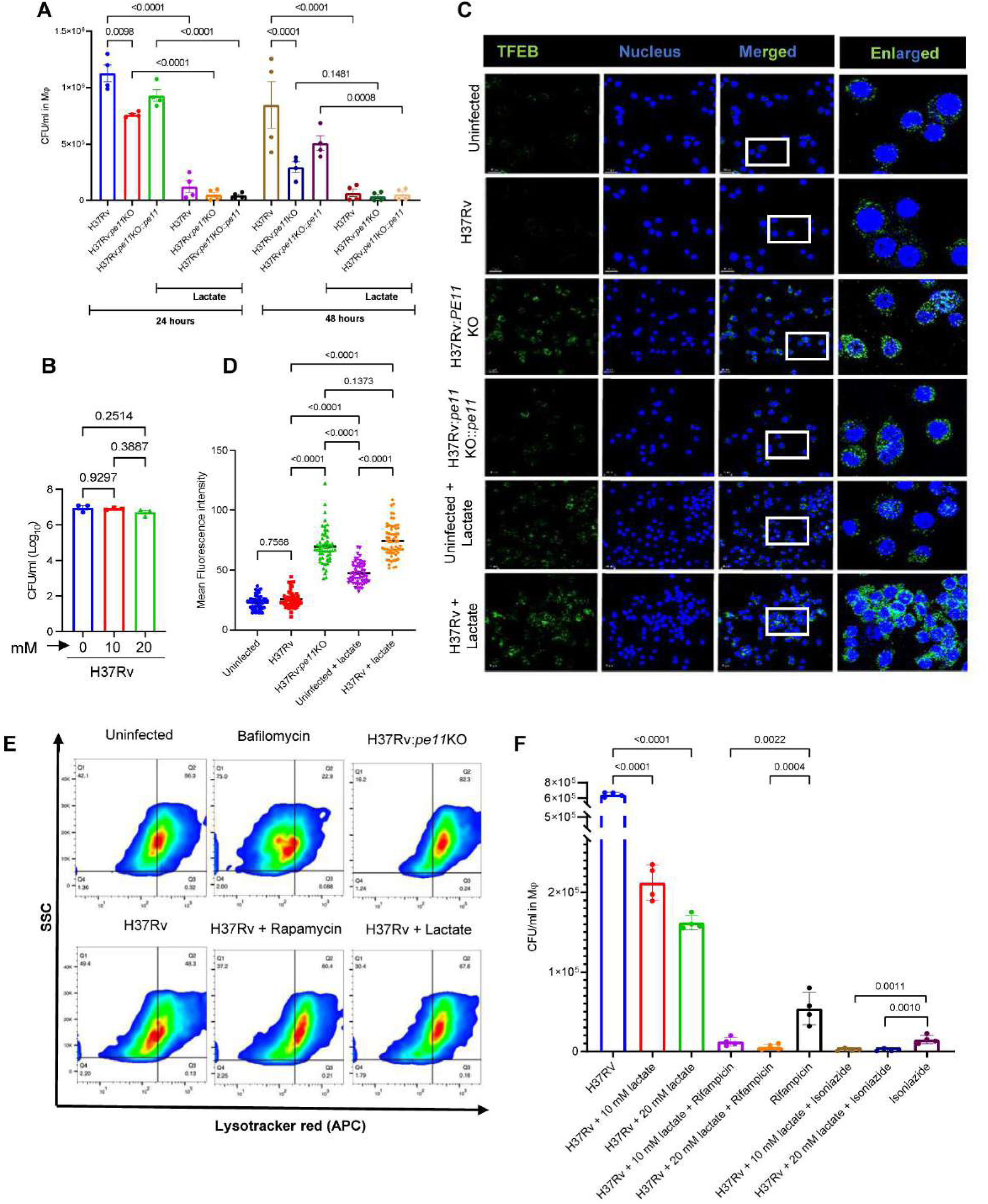
Lactate supplementation enhances host defence and promotes *M. tuberculosis* clearance. **(A)** Assay of intracellular mycobacterial survival in peritoneal macrophages infected with H37Rv, H37Rv:*pe11*KO, or *pe11*-complemented *pe11*KO strain (MOI 5) in the presence or absence of 10 mM lactate. Data presented as mean ± SEM (n = 4). **(B)** Mtb axenic culture were either left untreated or treated with lactate (10 mM or 20 mM) and bacterial survival assay was performed. Data represents mean ± SEM of 3 experiments. (**C**) Confocal images showing TFEB nuclear translocation in H37Rv-infected BMC2 macrophages in the absence or presence of 10 mM lactate (TFEB: green; DAPI: blue; scale bar: 10 μm). **(D)** Mean fluorescence intensity representing the levels of total TFEB. Data presented as mean ± SEM. **(E)** Measurement of lysosomal acidification by LysoTracker Red using flow cytometry in C57BL/6 macrophages infected with various *M. tuberculosis* strains in the presence or absence of lactate (10 mM). **(F)** Bacterial survival in macrophages treated with either lactate (10 mM) alone or in combination with anti-TB drugs, Rifampicin (1 μg/ml) or Isoniazid (1 μg/ml) at 24 hpi and 48 hpi. Statistical significance was determined by one-way ANOVA. Cells left uninfected were used as control in all the experiments.

## Discussion

The PE11 (LipX, Rv1196c) protein of *M. tuberculosis* is a cell wall localized esterase which has been found to significantly remodel mycobacterial cell wall and confer survival advantage to the bacteria under hostile intracellular environmental conditions (Singh et al., 2016, Rastogi et al., 2017, Dahiya et al., 2024). It is also a member of the unique PE/PPE family of proteins found only in Mycobacteria and were initially predicted to play a role in immune evasion by serving as decoy antigen and generation of antigenic variation (Akhter et al., 2012). Like many other PE/PPE proteins, PE11 appears to be pleiotropic in nature, which can cause cell wall remodelling by changing the composition of the cell wall lipids as well as plays a role in creating a pro-inflammatory milieu by triggering production of pro-inflammatory cytokines during infection (Singh et al., 2016; Rastogi et al., 2017). We observed that PE11 plays a key role in Mtb stress adaptation. The *pe11*-knockout strain exhibits reduced bacillary size and impaired acid tolerance, consistent with the upregulation of *pe11* transcript in acidic conditions. Bacterial morphology is linked to stress resistance and virulence (Payros et al., 2021; Vijay et al., 2017), emphasising that PE11 probably enables Mtb to maintain structural resilience under immune pressure. Furthermore, PE11 has been shown to play a role in inducing necrosis in macrophages (Deng et al., 2015). This prompted us to examine the broader impact of PE11 on host cellular processes during mycobacterial infection.

Accordingly, we examined the changes in the transcriptome in peritoneal macrophages infected with a wild type virulent laboratory strain H37Rv and compared with that of a *pe11*-deficient H37Rv strain (*pe11*KO). Gene ontology enrichment analyses indicated that several genes were differentially regulated in the presence of PE11, predominantly belong to the immune pathways. True to these observations, we found that mice infected with PE11-expressing strains exhibited marked neutrophilia and monocytosis indicating a role of PE11 in inducing inflammation during infection. In contrast, mice infected with *pe11*-KO H37Rv strain, showed attenuated inflammation and reduced pulmonary lesions. Additionally, increased infiltration of CD45^+^ immune cells in the lungs and elevated serum pro-inflammatory cytokines like TNF-α, IL-6 and IL-1β, in the presence of PE11, further emphasize its role in mediating inflammation corroborating previous observations using a surrogate *M. smegmatis* expressing PE11 (Singh et al., 2016). As alveolar macrophages provide an early niche for Mtb infection and dissemination (Cohen et al. 2018), we further analyzed the inflammatory cytokine profile of primary macrophages infected wild-type and *pe11*KO Mtb strain. The results demonstrate that PE11 promotes pro-inflammatory responses in macrophages. Such dysregulated immune response is critical for Mtb persistence, as excessive inflammation can promote granuloma necrosis and bacterial dissemination (Davis et al., 2009; Elkington et al., 2022).

A central finding of this study is the PE11’s ability to subvert host autophagy, a key defence mechanism against intracellular pathogens (Siqueira et al., 2018; Yuk et al., 2012). Analyses of the transcript profile from the macrophages infected with *pe11*KO strain, suggest that in the absence of PE11, many genes related to autophagy and lysosomal biogenesis are significantly upregulated when compared with that of macrophages infected with wild type strain harbouring functional *pe11* gene. Therefore, in the absence of PE11, important host protective mechanisms are not significantly compromised, but are suppressed in PE11-sufficient strains. Genes critical to autophagy and lysosomal biogenesis, including atg5, *becn1, fnip1*, *fnip2*, and TFEB, were found to be significantly downregulated in the presence of PE11. These genes are critical for host defence mechanisms like autophagy and lysosomal functions that control intracellular infections, further underscoring PE11’s role in tipping the balance of host responses to favour pathogen survival. Autophagy associated genes like *atg5* and *beclin1* are known to be crucial for clearance of infected mycobacteria (Golovkine et al., 2023; Deretic, 2014; Kang et al., 2011; Changotra et al., 2022). The levels of *atg5* and *becn1* mRNA levels were found to be lower both in macrophages and in mice lung tissue infected with PE11-sufficient strains. PE11-mediated suppression of autophagy was further confirmed by elevated accumulation of p62 and decreased levels of LC3B. Autophagy is known to be critical for degradation of protein/antigens by delivering them into lysosomes (Birgisdottir and Johansen, 2020). We found that in the presence of PE11, DQ-OVA degradation was hampered, suggesting that PE11 disrupts the autophagic flow needed for proteolytic degradation in lysosomes. Since lysosomal acidification is critical for the optimal function of the proteolytic enzymes (Mindell, 2012), we speculated that PE11 may be involved in interfering with lysosomal acidification. True to our expectation it was found that, macrophages infected with *pe11*-sufficient strains had lesser fluorescence compared to that of macrophages infected with *pe11*-deficient strains, when incubated with LysoTracker, a dye activated in the acidic environment of lysosomes. As V-ATPases are essential for lysosmal acidification (Mindell, 2012), on examination it was found that the expression of V-ATPase subunits *atp6v1h*, and *atp6v0e* are significantly downregulated in infected macrophages as well as in the lung of infected mice with *pe11*-sufficient Mtb strains as compared to those infected with *pe11*-deficient strains. In fact, our transcriptomic data showed significant inhibition of several V-ATPase subunits (atp6v_0_C, atp6v_0_e2, atp6v_0_d2, atp6v_1_h, and atp6 v_0_E). Given the established role of the V-ATPase complex in lysosome acidification and biogenesis (Perera and Zoncu, 2016; Bouché et al., 2016; Maxson et al., 2014), our data suggests that PE11 probably disrupts lysosomal acidification, and thus protects the infected mycobacteria from degradation in the lysosome. This observation is consistent with previous studies connecting lysosomal pH imbalance to pathogen persistence (Sachdeva and Sundarmurthy, 2020; Teixeira et al., 2023). An interesting aspect of our study is that, PE11 was found to affect host autophagy and lysosomal function mechanistically by regulating TFEB activity. TFEB is a key regulator of several genes associated with autophagy and lysosomal biogenesis, including *atg5*, *becn1*, and V-ATPase subunits (Sardiello et al., 2009; Settembre et al., 2011; Cortes et al., 2019; Song et al., 2021; Yang and Wang, 2021; Gros and Muller, 2023; Medina et al., 2011; Settembre et al., 2012). Compared to wild-type Mtb infection, *pe11*KO-infected macrophages and mouse lung tissues displayed significantly higher TFEB expression, at both mRNA and protein levels indicating a role of PE11 in actively suppressing TFEB during mycobacterial infection and thereby suppresses expression of lysosome acidification and autophagy-related genes, which are crucial for efficient lysosomal acidification and autophagy initiation (Napolitano and Ballabio, 2016; Sardiello et al., 2009; Settembre et al., 2011; Cortes et al., 2019; Song et al., 2021; Yang and Wang, 2021; Gros and Muller, 2023). Recently, Mtb was found to inhibit TFEB activity through other factors using various other mechanisms also (Zhao et al., 2024; Biswas et al., 2023). Interestingly, accumulation of and activation of TFEB is negatively regulated by a tumour suppressor FLCN (Paquette et al., 2021), which also plays a role in regulating cellular lactate levels by acting as an uncompetitive inhibitor of lactate dehydrogenase A (LDHA), a glycolytic enzyme, in addition to its canonical role in mTOR regulation (Woodford et al., 2021). Interestingly, lactate has been shown to stabilize TFEB through lactylation, a post-translational modification essential for maintaining TFEB stability and promoting autophagic processes (Huang et al., 2024). Lactylation of TFEB at ^91^Lysine residue prevents TFEB to interact with E3 ubiquitin ligase WWP2, thereby inhibiting its ubiquitination and subsequent proteasome-mediated degradation (Huang et al., 2024). Therefore, the reduced lactate levels observed in *pe11*-expressing strains may impair TFEB lactylation, leading to its proteolytic degradation and diminished activity. Therefore, these data suggest a novel role of PE11 as a virulence factor which promotes mycobacterial survival by targeting the FLCN-lactate-TFEB axis to inhibit lysosomal acidification and impair autophagy.

Consistent to our findings, reduced lactate production was observed in bone marrow-derived macrophages (BMDM) during Mtb infection compared to those infected with heat-killed *Mycobacterium* (Ó Maoldomhnaigh et al., 2021). Additionally, exogenous lactate treatment enhanced clearance of irradiated Mtb in human macrophages (Ó Maoldomhnaigh et al., 2021), similar to our study performed with live Mtb, Similar effect of lactate in clearing mycobacteria was further confirmed *in vivo* using a mice model of infection with *Msmeg*-PE11.

It appears that one of the mechanisms by which PE11 exerts its virulence is through reduction of cellular lactate levels which in turn affects TFEB stability and subsequently autophagy and related defensive mechanisms of the host. This prompted us to speculate that supplementation with extracellular lactate may stabilize TFEB activity which may subsequently improve lysosome function and autophagy and improved the host’s ability to clear mycobacterial infections. True to our expectations, we found that supplementation of lactate in macrophages restored nuclear accumulation of TFEB and increased lysosomal acidification, leading to enhanced Mtb clearance. Therefore, FLCN-lactate-TFEB axis appears to be the primary target of PE11 to subvert autophagy, where PE11 play a role in orchestrating both metabolic (lactate depletion) and transcriptional (suppression of TFEB expression) function to impair host defence mechanisms. Supplementation of lactate alone caused >80% reduction in intracellular Mtb survival in macrophages, while in combination with first-line antibiotics cleared >95% of the bacteria. These data highlight the therapeutic potential of lactate as a novel host-directed therapeutic (HDT) and a potential adjutant to circumvent drug resistance (Young et al., 2020; Tian et al., 2025). HDTs are recently gaining traction as promising adjunct strategies to enhance tuberculosis (TB) treatment outcomes (Maeurer et al., 2016; Young et al., 2020; Tian et al., 2025). Unlike conventional antibiotics that target *M. tuberculosis* directly, HDTs aim to bolster the host’s immune response. This approach offers potential benefits such as reduced treatment duration, lower risk of drug resistance, and minimized lung pathology. Autophagy is a prominent host defense mechanism against Mtb (Gutierrez et al., 2004; Levine et al., 2008; Deretic, 2014; Siqueira et al., 2018), and enhancing this pathway may significantly improve immune control of the host and thereby limit bacterial persistence.

In summary, we describe a novel mechanism by which Mtb PE11, as a virulence factor, orchestrates a multipronged disruption of the host protective immune mechanisms, by targeting autophagy, lysosomal acidification, and cellular metabolism. By interfering with the FLCN-TFEB signaling axis, PE11 impairs lysosomal acidification and autophagic flux, while concurrently reducing cellular lactate levels to hamper TFEB stability. These coordinated effects help to create an environment that favours *M. tuberculosis* survival within the macrophages. Most importantly, we have been able to uncover potentiality of lactate as a candidate for HDT against *M. tuberculosis.* Collectively, these findings highlight the mechanisms by which PE11 undermines the host defense mechanisms, thereby promoting bacterial persistence and virulence.

## Acknowledgements

We thank ABSL3/BSL3 facility of International Centre for Genetic Engineering and Biotechnology (ICGEB), New Delhi for providing access to the ABSL3/BSL3 facility for the work related to the use of the pathogenic strain of *Mycobacterium tuberculosis.* We thank Dr Dhiraj Kumar and his team members for their assistance during the mouse aerosol infection. We also thank Dr Sandeep Kumar Kotturu and Priyanka Kushwaha for kind help. The authors gratefully acknowledge the financial support by the Science and Engineering Research Board (SERB), Department of Science and Technology (DST), Government of India (JCB/2021/000035), Council of Scientific and Industrial Research (CSIR), Govt. of India (37WS(0020)/2023-24/EMR-II/ASPIRE), Department of Biotechnology (DBT), Govt of India (BT/PR51149/MED/29/1660/2023) to SM and SG, Indian Council of Medical Research (ICMR), Govt. of India (2021-10087/GTGE/ADHOC-BMS and IIRPSG-2024-01-01453) and a core grant from CDFD by DBT. VM was supported by the DST-INSPIRE fellowship, Govt. of India.

## Author contributions

P.D. and S.M. conceived, designed the experiments, and analyzed the data. P.D performed most of the experiments. M.K.B. was involved in analyses of RNA-Seq data and RT-PCR. A.S. helped in H37Rv culture experiments. S.G. helped in blood profiling. V.K.N. provided support and facility of M. tuberculosis work at the BSL3 facility of CCMB, Hyderabad. A.C. and S.S.K helped in lactate assay. P.D., and S.M. wrote the manuscript, S.G. and S.M. corrected and edited the manuscript and S.M. provided guidance and financial support.

## Figure Legends

**Supplemental Figure 1. Mice infected with *M. smegmatis* expressing Mtb PE11, exhibit neutrophilia, monocytosis and increased infiltration of CD45+ immune cells into the lungs.** C57BL/6 mice were infected with 50 × 10^6^ CFUs of either *Msmeg-*pVV or *Msmeg-*PE11 through the intraperitoneal route. Uninfected mice were used as healthy control. At day 3 post infection, blood samples were collected and analyzed for levels of **(A)** Neutrophils and **(B)** Monocytes. Data are representative of mean[±[SEM of 5 mice. **(C)** Also, lungs were harvested and paraffin sections were prepared and stained with haematoxylin and eosin. Scale bar of top panel is 500 µm and bottom panel is 100 µm. **(D)** Lung sections were probed with anti-CD45 antibody and counterstained with haematoxylin and lung tissue from uninfected mice was taken as healthy control. Arrows indicate sign of inflammation and immune cell accumulation. Photographs of representative sections were visualized at 20X magnification. Scale bar of top lane is 100 µm and bottom lane is enlarged image of the section.

**Supplementary Figure 2. Heat map of top 50 differentially expressed genes.** C57BL/6 mice peritoneal macrophages were infected (1:5 MOI, 12 hpi), RNA sequencing is carried out. Heat-map of DEGs (columns) across infected macrophages (Rows), clustered by expression patterns is presented. Color scale: log2 normalized counts.

**Supplementary Figure 3. PE11 suppresses autophagy and dampened lysosomal acidification during *M. smegmatis* infection.** Analysis of autophagy markers (LC3B and p62) in C57BL/6 peritoneal macrophages infected with *Msmeg-*pVV or *Msmeg-*PE11 at MOI 1:10 (6 hours post-infection (hpi)). **(A)** Representative Western blots of LC3B and p62. Quantification normalized to β-actin. Data show mean ± SEM of 3 experiments. **(B)** Confocal microscopy showing LC3B puncta formation in BMC2 macrophages infected as above (LC3B: red; DAPI: blue; scale bar: 10 μm).

**Supplementary Figure 4. Effect of lactate supplementation on cell cytotoxicity (A, B)** Peritoneal macrophages from C57BL/6 mice were treated with various concentrations of lactate for 24 hours (A) and 48 hours (B) and cytotoxicity assay was performed by MTT assay. Data (mean ± SEM of 3 different experiments) are shown as percentage viability of the untreated cultures.

**Supplementary Figure 5. Lactic acid inhibits mycobacterial growth in mice. (A)** C57BL/6 peritoneal macrophages infected with H37Rv (MOI 5, 12 hpi). And representative images of live macrophages with or without lactate (10 mM) are represented in the panel. Statistical significance was calculated using one-way ANOVA.

## References

1. Ahmed A, Das A, Mukhopadhyay S. Immunoregulatory functions and expression patterns of PE/PPE family members: Roles in pathogenicity and impact on anti-tuberculosis vaccine and drug design. IUBMB Life. 2015 Jun;67(6):414–27. doi: 10.1002/iub.1387. Epub 2015 Jun 24. PMID: 26104967.

2. Akhter Y, Ehebauer MT, Mukhopadhyay S, Hasnain SE. The PE/PPE multigene family codes for virulence factors and is a possible source of mycobacterial antigenic variation: perhaps more? Biochimie. 2012 Jan;94(1):110–6. doi: 10.1016/j.biochi.2011.09.026. Epub 2011 Oct 12. PMID: 22005451.

3. Ayyappan JP, Ganapathi U, Lizardo K, Vinnard C, Subbian S, Perlin DS, Nagajyothi JF. Adipose Tissue Regulates Pulmonary Pathology during TB Infection. mBio. 2019 Apr 16;10(2):e02771–18. doi: 10.1128/mBio.02771-18. PMID: 30992360; PMCID: PMC6469978.

4. Ballabio A, Bonifacino JS. Lysosomes as dynamic regulators of cell and organismal homeostasis. Nat Rev Mol Cell Biol. 2020 Feb;21(2):101–118. doi: 10.1038/s41580-019-0185-4. Epub 2019 Nov 25. PMID: 31768005.

5. Betts JC, Lukey PT, Robb LC, McAdam RA, Duncan K. Evaluation of a nutrient starvation model of Mycobacterium tuberculosis persistence by gene and protein expression profiling. Mol Microbiol. 2002 Feb;43(3):717–31. doi: 10.1046/j.1365-2958.2002.02779.x. PMID: 11929527.

6. Birgisdottir ÅB, Johansen T. Autophagy and endocytosis - interconnections and interdependencies. J Cell Sci. 2020 May 22;133(10):jcs228114. doi: 10.1242/jcs.228114. PMID: 32501285.

7. Bisht MK, Pal R, Dahiya P, Naz S, Sanyal P, Nandicoori VK, Ghosh S, Mukhopadhyay S. The PPE2 protein of *Mycobacterium tuberculosis* is secreted during infection and facilitates mycobacterial survival inside the host. Tuberculosis (Edinb). 2023 Dec;143:102421. doi: 10.1016/j.tube.2023.102421. Epub 2023 Oct 12. PMID: 37879126.

8. Biswas VK, Sen K, Ahad A, Ghosh A, Verma S, Pati R, Prusty S, Nayak SP, Podder S, Kumar D, Gupta B, Raghav SK. NCoR1 controls *Mycobacterium tuberculosis* growth in myeloid cells by regulating the AMPK-mTOR-TFEB axis. PLoS Biol. 2023 Aug 17;21(8):e3002231. doi: 10.1371/journal.pbio.3002231. PMID: 37590294; PMCID: PMC10465006.

9. Bouché V, Espinosa AP, Leone L, Sardiello M, Ballabio A, Botas J. Drosophila Mitf regulates the V-ATPase and the lysosomal-autophagic pathway. Autophagy. 2016;12(3):484–98. doi: 10.1080/15548627.2015.1134081. PMID: 26761346; PMCID: PMC4835958.

10. Bradfute SB, Castillo EF, Arko-Mensah J, Chauhan S, Jiang S, Mandell M, Deretic V. Autophagy as an immune effector against tuberculosis. Curr Opin Microbiol. 2013 Jun;16(3):355–65. doi: 10.1016/j.mib.2013.05.003. Epub 2013 Jun 18. PMID: 23790398; PMCID: PMC3742717.

11. Brennan PJ, Nikaido H. The envelope of mycobacteria. Annu Rev Biochem. 1995;64:29–63. doi: 10.1146/annurev.bi.64.070195.000333. PMID: 7574484.

12. Castillo EF, Dekonenko A, Arko-Mensah J, Mandell MA, Dupont N, Jiang S, Delgado-Vargas M, Timmins GS, Bhattacharya D, Yang H, Hutt J, Lyons CR, Dobos KM, Deretic V. Autophagy protects against active tuberculosis by suppressing bacterial burden and inflammation. Proc Natl Acad Sci U S A. 2012 Nov 13;109(46):E3168–76. doi: 10.1073/pnas.1210500109. PMID: 23093667; PMCID: PMC3503152.

13. Chandra P, Grigsby SJ, Philips JA. Immune evasion and provocation by *Mycobacterium tuberculosis*. Nat Rev Microbiol. 2022 Dec;20(12):750–766. doi: 10.1038/s41579-022-00763-4.

14. Changotra H, Kaur S, Yadav SS, Gupta GL, Parkash J, Duseja A. ATG5: A central autophagy regulator implicated in various human diseases. Cell Biochem Funct. 2022 Oct;40(7):650–667. doi: 10.1002/cbf.3740. Epub 2022 Sep 5. PMID: 36062813.

15. Chen YY, Liu CX, Liu HX, Wen SY. The emerging roles of vacuolar-type ATPase-dependent lysosomal acidification in cardiovascular disease. Biomolecules. 2025 Apr 3;15(4):525. doi: 10.3390/biom15040525. PMID: 40305271; PMCID: PMC12024769.

16. Cohen SB, Gern BH, Delahaye JL, Adams KN, Plumlee CR, Winkler JK, Sherman DR, Gerner MY, Urdahl KB. Alveolar macrophages provide an early *Mycobacterium tuberculosis* niche and initiate dissemination. Cell Host Microbe. 2018 Sep 12;24(3):439–446.e4. doi: 10.1016/j.chom.2018.08.001. Epub 2018 Aug 23. PMID: 30146391; PMCID: PMC6152889.

17. Colacurcio DJ, Nixon RA. Disorders of lysosomal acidification-The emerging role of v-ATPase in aging and neurodegenerative disease. Ageing Res Rev. 2016 Dec;32:75–88. doi: 10.1016/j.arr.2016.05.004. Epub 2016 May 16. PMID: 27197071; PMCID: PMC5112157.

18. Cortes CJ, La Spada AR. TFEB dysregulation as a driver of autophagy dysfunction in neurodegenerative disease: Molecular mechanisms, cellular processes, and emerging therapeutic opportunities. Neurobiol Dis. 2019 Feb;122:83–93. doi: 10.1016/j.nbd.2018.05.012. Epub 2018 May 28. PMID: 29852219; PMCID: PMC6291370.

19. Cronan MR. In the Thick of It: Formation of the tuberculous granuloma and its effects on host and therapeutic responses. Front Immunol. 2022 Mar 7;13:820134. doi: 10.3389/fimmu.2022.820134. PMID: 35320930; PMCID: PMC8934850.

20. Dahiya P, Banerjee A, Saha A, Nandicoori VK, Ghosh S, Mukhopadhyay S. Structure-function relationship of PE11 esterase of *Mycobacterium tuberculosis* with respect to its role in virulence. Biochem Biophys Res Commun. 2024 Dec 20;739:150927. doi: 10.1016/j.bbrc.2024.150927. Epub 2024 Nov 10. PMID: 39541926.

21. Davis JM, Ramakrishnan L. The role of the granuloma in expansion and dissemination of early tuberculous infection. Cell. 2009 Jan 9;136(1):37–49. doi: 10.1016/j.cell.2008.11.014. PMID: 19135887; PMCID: PMC3134310.

22. Daniel J, Maamar H, Deb C, Sirakova TD, Kolattukudy PE. *Mycobacterium tuberculosis* uses host triacylglycerol to accumulate lipid droplets and acquires a dormancy-like phenotype in lipid-loaded macrophages. PLoS Pathog. 2011 Jun;7(6):e1002093. doi: 10.1371/journal.ppat.1002093. Epub 2011 Jun 23. PMID: 21731490; PMCID: PMC3121879.

23. Deng W, Zeng J, Xiang X, Li P, Xie J. PE11 (Rv1169c) selectively alters fatty acid components of *Mycobacterium smegmatis* and host cell interleukin-6 level accompanied with cell death. Front Microbiol. 2015 Jun 23;6:613. doi: 10.3389/fmicb.2015.00613. PMID: 26157429; PMCID: PMC4477156.

24. Deretic V. Autophagy in tuberculosis. Cold Spring Harb Perspect Med. 2014 Aug 28;4(11):a018481. doi: 10.1101/cshperspect.a018481. PMID: 25167980; PMCID: PMC4208715.

25. El-Houjeiri L, Possik E, Vijayaraghavan T, Paquette M, Martina JA, Kazan JM, Ma EH, Jones R, Blanchette P, Puertollano R, Pause A. The transcription factors TFEB and TFE3 link the FLCN-AMPK signaling axis to innate immune response and pathogen resistance. Cell Rep. 2019 Mar 26;26(13):3613–3628.e6. doi: 10.1016/j.celrep.2019.02.102. PMID: 30917316; PMCID: PMC7457953.

26. Elkington P, Polak ME, Reichmann MT, Leslie A. Understanding the tuberculosis granuloma: the matrix revolutions. Trends Mol Med. 2022 Feb;28(2):143–154. doi: 10.1016/j.molmed.2021.11.004. Epub 2021 Dec 15. PMID: 34922835; PMCID: PMC8673590.

27. Etna MP, Giacomini E, Severa M, Coccia EM. Pro- and anti-inflammatory cytokines in tuberculosis: a two-edged sword in TB pathogenesis. Semin Immunol. 2014 Dec;26(6):543–51. Doi: 10.1016/j.smim.2014.09.011. Epub 2014 Nov 8. PMID: 25453229.

28. Fisher MA, Plikaytis BB, Shinnick TM. Microarray analysis of the *Mycobacterium tuberculosis* transcriptional response to the acidic conditions found in phagosomes. J Bacteriol. 2002 Jul;184(14):4025–32. doi: 10.1128/JB.184.14.4025-4032.2002. PMID: 12081975; PMCID: PMC135184.

29. Fontán PA, Voskuil MI, Gomez M, Tan D, Pardini M, Manganelli R, Fattorini L, Schoolnik GK, Smith I. The *Mycobacterium tuberculosis* sigma factor sigmaB is required for full response to cell envelope stress and hypoxia in vitro, but it is dispensable for in vivo growth. J Bacteriol. 2009 Sep;191(18):5628–33. doi: 10.1128/JB.00510-09. Epub 2009 Jul 10. PMID: 19592585; PMCID: PMC2737972.

30. Global tuberculosis report 2024. Geneva: World Health Organization; 2024. Licence: CC BY-NC-SA 3.0 IGO.

31. Golovkine GR, Roberts AW, Morrison HM, Rivera-Lugo R, McCall RM, Nilsson H, Garelis NE, Repasy T, Cronce M, Budzik J, Van Dis E, Popov LM, Mitchell G, Zalpuri R, Jorgens D, Cox JS. Autophagy restricts *Mycobacterium tuberculosis* during acute infection in mice. Nat Microbiol. 2023 May;8(5):819–832. doi: 10.1038/s41564-023-01354-6. Epub 2023 Apr 10. Erratum in: Nat Microbiol. 2024 Jul;9(7):1899. doi: 10.1038/s41564-024-01605-0. PMID: 37037941; PMCID: PMC11027733.

32. Gray MA, Choy CH, Dayam RM, Ospina-Escobar E, Somerville A, Xiao X, Ferguson SM, Botelho RJ. Phagocytosis enhances lysosomal and bactericidal properties by activating the transcription factor TFEB. Curr Biol. 2016 Aug 8;26(15):1955–1964. doi: 10.1016/j.cub.2016.05.070. Epub 2016 Jul 7. PMID: 27397893; PMCID: PMC5453720.

33. Gros F, Muller S. The role of lysosomes in metabolic and autoimmune diseases. Nat Rev Nephrol. 2023 Jun;19(6):366–383. doi: 10.1038/s41581-023-00692-2. Epub 2023 Mar 9. PMID: 36894628.

34. Gutierrez MG, Master SS, Singh SB, Taylor GA, Colombo MI, Deretic V. Autophagy is a defense mechanism inhibiting BCG and *Mycobacterium tuberculosis* survival in infected macrophages. Cell. 2004 Dec 17;119(6):753–66. doi: 10.1016/j.cell.2004.11.038. PMID: 15607973.

35. Houben RM, Dodd PJ. The global burden of latent tuberculosis infection: A re-estimation using mathematical modeling. PLoS Med. 2016 Oct 25;13(10):e1002152. doi: 10.1371/journal.pmed.1002152. PMID: 27780211; PMCID: PMC5079585.

36. Huang Y, Luo G, Peng K, Song Y, Wang Y, Zhang H, Li J, Qiu X, Pu M, Liu X, Peng C, Neculai D, Sun Q, Zhou T, Huang P, Liu W. Lactylation stabilizes TFEB to elevate autophagy and lysosomal activity. J Cell Biol. 2024 Nov 4;223(11):e202308099. doi: 10.1083/jcb.202308099. Epub 2024 Aug 28. PMID: 39196068; PMCID: PMC11354204.

37. Jain N, Kalam H, Singh L, Sharma V, Kedia S, Das P, Ahuja V, Kumar D. Mesenchymal stem cells offer a drug-tolerant and immune-privileged niche to *Mycobacterium tuberculosis*. Nat Commun. 2020 Jun 16;11(1):3062. doi: 10.1038/s41467-020-16877-3. PMID: 32546788; PMCID: PMC7297998.

38. Kalscheuer R, Palacios A, Anso I, Cifuente J, Anguita J, Jacobs WR Jr, Guerin ME, Prados-Rosales R. The *Mycobacterium tuberculosis* capsule: a cell structure with key implications in pathogenesis. Biochem J. 2019 Jul 18;476(14):1995–2016. doi: 10.1042/BCJ20190324. PMID: 31320388; PMCID: PMC6698057.

39. Kang R, Zeh HJ, Lotze MT, Tang D. The Beclin 1 network regulates autophagy and apoptosis. Cell Death Differ. 2011 Apr;18(4):571–80. doi: 10.1038/cdd.2010.191.

40. Kaufmann SHE, Dorhoi A, Hotchkiss RS, Bartenschlager R. Host-directed therapies for bacterial and viral infections. Nat Rev Drug Discov. 2018 Jan;17(1):35–56. doi: 10.1038/nrd.2017.162. Epub 2017 Sep 22. PMID: 28935918; PMCID: PMC7097079.

41. Kim YS, Silwal P, Kim SY, Yoshimori T, Jo EK. Autophagy-activating strategies to promote innate defense against mycobacteria. Exp Mol Med. 2019 Dec 11;51(12):1–10. doi: 10.1038/s12276-019-0290-7. PMID: 31827065; PMCID: PMC6906292.

42. Kimmey JM, Huynh JP, Weiss LA, Park S, Kambal A, Debnath J, Virgin HW, Stallings CL. Unique role for ATG5 in neutrophil-mediated immunopathology during *M. tuberculosis* infection. Nature. 2015 Dec 24;528(7583):565–9. doi: 10.1038/nature16451. Epub 2015 Dec 9. PMID: 26649827; PMCID: PMC4842313.

43. Kimmey JM, Stallings CL. Bacterial Pathogens versus Autophagy: Implications for Therapeutic Interventions. Trends Mol Med. 2016 Dec;22(12):1060–1076. doi: 10.1016/j.molmed.2016.10.008. Epub 2016 Nov 17. PMID: 27866924; PMCID: PMC5215815.

44. Lawrence RE, Zoncu R. The lysosome as a cellular centre for signalling, metabolism and quality control. Nat Cell Biol. 2019 Feb;21(2):133–142. doi: 10.1038/s41556-018-0244-7. Epub 2019 Jan 2. PMID: 30602725.

45. Levine B, Kroemer G. Autophagy in the pathogenesis of disease. Cell. 2008 Jan 11;132(1):27–42. doi: 10.1016/j.cell.2007.12.018. PMID: 18191218; PMCID: PMC2696814.

46. Lillebaek T, Dirksen A, Baess I, Strunge B, Thomsen VØ, Andersen AB. Molecular evidence of endogenous reactivation of *Mycobacterium tuberculosis* after 33 years of latent infection. J Infect Dis. 2002 Feb 1;185(3):401–4. doi: 10.1086/338342. Epub 2002 Jan 17. PMID: 11807725.

47. Lin H, Xing J, Wang H, Wang S, Fang R, Li X, Li Z, Song N. Roles of Lipolytic enzymes in *Mycobacterium tuberculosis* pathogenesis. Front Microbiol. 2024 Jan 29;15:1329715. doi: 10.3389/fmicb.2024.1329715. PMID: 38357346; PMCID: PMC10865251

48. Maeurer M, Rao M, Zumla A. Host-directed therapies for antimicrobial resistant respiratory tract infections. Curr Opin Pulm Med. 2016 May;22(3):203–11. doi: 10.1097/MCP.0000000000000271. PMID: 26989822.

49. Maxson ME, Grinstein S. The vacuolar-type H[-ATPase at a glance - more than a proton pump. J Cell Sci. 2014 Dec 1;127(Pt 23):4987–93. doi: 10.1242/jcs.158550. PMID: 25453113.

50. McCaffrey EF, Donato M, Keren L, Chen Z, Delmastro A, Fitzpatrick MB, Gupta S, Greenwald NF, Baranski A, Graf W, Kumar R, Bosse M, Fullaway CC, Ramdial PK, Forgó E, Jojic V, Van Valen D, Mehra S, Khader SA, Bendall SC, van de Rijn M, Kalman D, Kaushal D, Hunter RL, Banaei N, Steyn AJC, Khatri P, Angelo M. Author Correction: The immunoregulatory landscape of human tuberculosis granulomas. Nat Immunol. 2022 May;23(5):814. doi: 10.1038/s41590-022-01178-2. Erratum for: Nat Immunol. 2022 Feb;23(2):318-329. doi: 10.1038/s41590-021-01121-x. PMID: 35277696; PMCID: PMC9098386.

51. Medina DL, Fraldi A, Bouche V, Annunziata F, Mansueto G, Spampanato C, Puri C, Pignata A, Martina JA, Sardiello M, Palmieri M, Polishchuk R, Puertollano R, Ballabio A. Transcriptional activation of lysosomal exocytosis promotes cellular clearance. Dev Cell. 2011 Sep 13;21(3):421–30. doi: 10.1016/j.devcel.2011.07.016. Epub 2011 Sep 1. PMID: 21889421; PMCID: PMC3173716.

52. Mindell JA. Lysosomal acidification mechanisms. Annu Rev Physiol. 2012;74:69-86. doi: 10.1146/annurev-physiol-012110-142317. PMID: 22335796.

53. Mizushima N, Yoshimori T, Levine B. Methods in mammalian autophagy research. Cell. 2010 Feb 5;140(3):313–26. doi: 10.1016/j.cell.2010.01.028. PMID: 20144757; PMCID: PMC2852113.

54. Mukhopadhyay S, Balaji KN. The PE and PPE proteins of *Mycobacterium tuberculosis*. Tuberculosis (Edinb). 2011 Sep;91(5):441–7. doi: 10.1016/j.tube.2011.04.004. Epub 2011 May 6. PMID: 21527209.

55. Napolitano G, Ballabio A. TFEB at a glance. J Cell Sci. 2016 Jul 1;129(13):2475–81. doi: 10.1242/jcs.146365. Epub 2016 Jun 1. PMID: 27252382; PMCID: PMC4958300.

56. Napolitano G, Di Malta C, Ballabio A. Non-canonical mTORC1 signaling at the lysosome. Trends Cell Biol. 2022 Nov;32(11):920–931. doi: 10.1016/j.tcb.2022.04.012. Epub 2022 May 30. PMID: 35654731.

57. Neyrolles O, Hernández-Pando R, Pietri-Rouxel F, Fornès P, Tailleux L, Barrios Payán JA, Pivert E, Bordat Y, Aguilar D, Prévost MC, Petit C, Gicquel B. Is adipose tissue a place for *Mycobacterium tuberculosis* persistence? PLoS One. 2006 Dec 20;1(1):e43. doi: 10.1371/journal.pone.0000043. PMID: 17183672; PMCID: PMC1762355.

58. Ó Maoldomhnaigh C, Cox DJ, Phelan JJ, Mitermite M, Murphy DM, Leisching G, Thong L, O’Leary SM, Gogan KM, McQuaid K, Coleman AM, Gordon SV, Basdeo SA, Keane J. Lactate alters metabolism in human macrophages and improves their ability to kill *Mycobacterium tuberculosis*. Front Immunol. 2021 Oct 6;12:663695. doi: 10.3389/fimmu.2021.663695. PMID: 34691015; PMCID: PMC8526932.

59. Øynebråten I. Involvement of autophagy in MHC class I antigen presentation. Scand J Immunol. 2020 Nov;92(5):e12978. doi: 10.1111/sji.12978. Epub 2020 Oct 19. PMID: 32969499; PMCID: PMC7685157.

60. Paquette M, Yan M, Ramírez-Reyes JMJ, El-Houjeiri L, Biondini M, Dufour CR, Jeong H, Pacis A, Giguère V, Estall JL, Siegel PM, Audet-Walsh É, Pause A. Loss of hepatic Flcn protects against fibrosis and inflammation by activating autophagy pathways. Sci Rep. 2021 Oct 28;11(1):21268. doi: 10.1038/s41598-021-99958-7. PMID: 34711912; PMCID: PMC8553785.

61. Payros D, Alonso H, Malaga W, Volle A, Mazères S, Déjean S, Valière S, Moreau F, Balor S, Stella A, Combes-Soia L, Burlet-Schiltz O, Bouchez O, Nigou J, Astarie-Dequeker C, Guilhot C. Rv0180c contributes to *Mycobacterium tuberculosis* cell shape and to infectivity in mice and macrophages. PLoS Pathog. 2021 Nov 1;17(11):e1010020. doi: 10.1371/journal.ppat.1010020. PMID: 34724002; PMCID: PMC8584747.

62. Perera RM, Zoncu R. The lysosome as a regulatory hub. Annu Rev Cell Dev Biol. 2016 Oct 6;32:223–253. doi: 10.1146/annurev-cellbio-111315-125125. Epub 2016 Aug 3. PMID: 27501449; PMCID: PMC9345128.

63. Plumlee CR, Duffy FJ, Gern BH, Delahaye JL, Cohen SB, Stoltzfus CR, Rustad TR, Hansen SG, Axthelm MK, Picker LJ, Aitchison JD, Sherman DR, Ganusov VV, Gerner MY, Zak DE, Urdahl KB. Ultra-low dose aerosol infection of mice with *Mycobacterium tuberculosis* more closely models human tuberculosis. Cell Host Microbe. 2021 Jan 13;29(1):68–82.e5. doi: 10.1016/j.chom.2020.10.003. Epub 2020 Nov 2. PMID: 33142108; PMCID: PMC7854984.

64. Rachman H, Strong M, Ulrichs T, Grode L, Schuchhardt J, Mollenkopf H, Kosmiadi GA, Eisenberg D, Kaufmann SH. Unique transcriptome signature of *Mycobacterium tuberculosis* in pulmonary tuberculosis. Infect Immun. 2006 Feb;74(2):1233–42. doi: 10.1128/IAI.74.2.1233-1242.2006. PMID: 16428773; PMCID: PMC1360294.

65. Rai R, Singh V, Mathew BJ, Singh AK, Chaurasiya SK. Mycobacterial response to an acidic environment: protective mechanisms. Pathog Dis. 2022 Oct 3;80(1):ftac032. doi: 10.1093/femspd/ftac032. PMID: 35953394.

66. Rajendran A, Soory A, Khandelwal N, Ratnaparkhi G, Kamat SS. A multi-omics analysis reveals that the lysine deacetylase ABHD14B influences glucose metabolism in mammals. J Biol Chem. 2022 Jul;298(7):102128. doi: 10.1016/j.jbc.2022.102128. Epub 2022 Jun 11. PMID: 35700823; PMCID: PMC9270251.

67. Rastogi S, Singh AK, Pant G, Mitra K, Sashidhara KV, Krishnan MY. Down-regulation of PE11, a cell wall associated esterase, enhances the biofilm growth of *Mycobacterium tuberculosis* and reduces cell wall virulence lipid levels. Microbiology (Reading). 2017 Jan;163(1):52–61. doi: 10.1099/mic.0.000417. PMID: 28198348.

68. Rubin EJ. The granuloma in tuberculosis--friend or foe? N Engl J Med. 2009 Jun 4;360(23):2471–3. doi: 10.1056/NEJMcibr0902539. PMID: 19494225.

69. Sachdeva K, Sundaramurthy V. The Interplay of Host Lysosomes and intracellular pathogens. Front Cell Infect Microbiol. 2020 Nov 20;10:595502. doi: 10.3389/fcimb.2020.595502. PMID: 33330138; PMCID: PMC7714789.

70. Saftig P, Klumperman J. Lysosome biogenesis and lysosomal membrane proteins: trafficking meets function. Nat Rev Mol Cell Biol. 2009 Sep;10(9):623–35. doi: 10.1038/nrm2745. Epub 2009 Aug 12. PMID: 19672277.

71. Sampson SL. Mycobacterial PE/PPE proteins at the host-pathogen interface. Clin Dev Immunol. 2011;2011:497203. doi: 10.1155/2011/497203. Epub 2011 Jan 26. PMID: 21318182; PMCID: PMC3034920.

72. Sapkota A, Park EJ, Kim YJ, Heo JB, Nguyen TQ, Heo BE, Kim JK, Lee SH, Kim SI, Choi YJ, Roh T, Jeon SM, Jang M, Heo HJ, Whang J, Paik S, Yuk JM, Kim JM, Song GY, Jang J, Jo EK. The autophagy-targeting compound V46 enhances antimicrobial responses to *Mycobacteroides abscessus* by activating transcription factor EB. Biomed Pharmacother. 2024 Oct;179:117313. doi: 10.1016/j.biopha.2024.117313. Epub 2024 Aug 20. PMID: 39167844.

73. Sardiello M, Palmieri M, di Ronza A, Medina DL, Valenza M, Gennarino VA, Di Malta C, Donaudy F, Embrione V, Polishchuk RS, Banfi S, Parenti G, Cattaneo E, Ballabio A. A gene network regulating lysosomal biogenesis and function. Science. 2009 Jul 24;325(5939):473-7. doi: 10.1126/science.1174447. Epub 2009 Jun 25. PMID: 19556463.

74. Schnappinger D, Ehrt S, Voskuil MI, Liu Y, Mangan JA, Monahan IM, Dolganov G, Efron B, Butcher PD, Nathan C, Schoolnik GK. Transcriptional adaptation of *Mycobacterium tuberculosis* within macrophages: Insights into the phagosomal environment. J Exp Med. 2003 Sep 1;198(5):693–704. doi: 10.1084/jem.20030846. PMID: 12953091; PMCID: PMC2194186.

75. Settembre C, Di Malta C, Polito VA, Garcia Arencibia M, Vetrini F, Erdin S, Erdin SU, Huynh T, Medina D, Colella P, Sardiello M, Rubinsztein DC, Ballabio A. TFEB links autophagy to lysosomal biogenesis. Science. 2011 Jun 17;332(6036):1429-33. doi: 10.1126/science.1204592. Epub 2011 May 26. PMID: 21617040; PMCID: PMC3638014.

76. Settembre C, Zoncu R, Medina DL, Vetrini F, Erdin S, Erdin S, Huynh T, Ferron M, Karsenty G, Vellard MC, Facchinetti V, Sabatini DM, Ballabio A. A lysosome-to-nucleus signalling mechanism senses and regulates the lysosome via mTOR and TFEB. EMBO J. 2012 Mar 7;31(5):1095–108. doi: 10.1038/emboj.2012.32. Epub 2012 Feb 17. PMID: 22343943; PMCID: PMC3298007.

77. Singh P, Rao RN, Reddy JR, Prasad RB, Kotturu SK, Ghosh S, Mukhopadhyay S. PE11, a PE/PPE family protein of *Mycobacterium tuberculosis* is involved in cell wall remodeling and virulence. Sci Rep. 2016 Feb 23;6:21624. doi: 10.1038/srep21624. PMID: 26902658; PMCID: PMC4763214

78. Siqueira MDS, Ribeiro RM, Travassos LH. Autophagy and its interaction with intracellular bacterial pathogens. Front Immunol. 2018 May 23;9:935. doi: 10.3389/fimmu.2018.00935. PMID: 29875765; PMCID: PMC5974045.

79. Song Q, Meng B, Xu H, Mao Z. The emerging roles of vacuolar-type ATPase-dependent Lysosomal acidification in neurodegenerative diseases. Transl Neurodegener. 2020 May 11;9(1):17. doi: 10.1186/s40035-020-00196-0. PMID: 32393395; PMCID: PMC7212675.

80. Song TT, Cai RS, Hu R, Xu YS, Qi BN, Xiong YA. The important role of TFEB in autophagy-lysosomal pathway and autophagy-related diseases: a systematic review. Eur Rev Med Pharmacol Sci. 2021 Feb;25(3):1641–1649. doi: 10.26355/eurrev_202102_24875. PMID: 33629334.

81. Sun X, Shu Y, Yan P, Huang H, Gao R, Xu M, Lu L, Tian J, Huang D, Zhang J. Transcriptome profiling analysis reveals that ATP6V0E2 is involved in the lysosomal activation by anlotinib. Cell Death Dis. 2020 Aug 24;11(8):702. doi: 10.1038/s41419-020-02904-0. PMID: 32839434; PMCID: PMC7445181.

82. Teixeira SC, Teixeira TL, Tavares PCB, Alves RN, da Silva AA, Borges BC, Martins FA, Dos Santos MA, de Castilhos P, E Silva Brígido RT, Notário AFO, Silveira ACA, da Silva CV. Subversion strategies of lysosomal killing by intracellular pathogens. Microbiol Res. 2023 Dec;277:127503. doi: 10.1016/j.micres.2023.127503. Epub 2023 Sep 22. PMID: 37748260.

83. Tian N, Chu H, Li Q, Sun H, Zhang J, Chu N, Sun Z. Host-directed therapy for tuberculosis. Eur J Med Res. 2025 Apr 11;30(1):267. doi: 10.1186/s40001-025-02443-4. PMID: 40211397; PMCID: PMC11987284.

84. Typas D. Autophagy counteracts *Mycobacterium tuberculosis* infection at early stages. Nat Struct Mol Biol. 2023 Jun;30(6):720. doi: 10.1038/s41594-023-01024-5. PMID: 37336992.

85. Velayati AA, Farnia P, Ibrahim TA, Haroun RZ, Kuan HO, Ghanavi J, Farnia P, Kabarei AN, Tabarsi P, Omar AR, Varahram M, Masjedi MR. Differences in cell wall thickness between resistant and nonresistant strains of *Mycobacterium tuberculosis*: using transmission electron microscopy. Chemotherapy. 2009;55(5):303–7. doi: 10.1159/000226425. Epub 2009 Jun 26. PMID: 19556787.

86. Vijay S, Nair RR, Sharan D, Jakkala K, Mukkayyan N, Swaminath S, Pradhan A, Joshi NV, Ajitkumar P. Mycobacterial cultures contain cell size and density specific sub-populations of cells with significant differential susceptibility to antibiotics, oxidative and nitrite stress. Front Microbiol. 2017 Mar 21;8:463. doi: 10.3389/fmicb.2017.00463. PMID: 28377757; PMCID: PMC5359288.

87. Voskuil MI, Visconti KC, Schoolnik GK. *Mycobacterium tuberculosis* gene expression during adaptation to stationary phase and low-oxygen dormancy. Tuberculosis (Edinb). 2004;84(3-4):218–27. doi: 10.1016/j.tube.2004.02.003. PMID: 15207491.

88. Voskuil MI, Bartek IL, Visconti K, Schoolnik GK. The response of *Mycobacterium tuberculosis* to reactive oxygen and nitrogen species. Front Microbiol. 2011 May 13;2:105. doi: 10.3389/fmicb.2011.00105. PMID: 21734908; PMCID: PMC3119406.

89. Woodford MR, Baker-Williams AJ, Sager RA, Backe SJ, Blanden AR, Hashmi F, Kancherla P, Gori A, Loiselle DR, Castelli M, Serapian SA, Colombo G, Haystead TA, Jensen SM, Stetler-Stevenson WG, Loh SN, Schmidt LS, Linehan WM, Bah A, Bourboulia D, Bratslavsky G, Mollapour M. The tumor suppressor folliculin inhibits lactate dehydrogenase A and regulates the Warburg effect. Nat Struct Mol Biol. 2021 Aug;28(8):662–670. doi: 10.1038/s41594-021-00633-2. Epub 2021 Aug 11. PMID: 34381247; PMCID: PMC9278990.

90. Yambire KF, Rostosky C, Watanabe T, Pacheu-Grau D, Torres-Odio S, Sanchez-Guerrero A, Senderovich O, Meyron-Holtz EG, Milosevic I, Frahm J, West AP, Raimundo N. Impaired lysosomal acidification triggers iron deficiency and inflammation in vivo. Elife. 2019 Dec 3;8:e51031. doi: 10.7554/eLife.51031. PMID: 31793879; PMCID: PMC6917501.

91. Yang DC, Blair KM, Salama NR. Staying in shape: the impact of cell shape on bacterial survival in diverse environments. Microbiol Mol Biol Rev. 2016 Feb 10;80(1):187–203. doi: 10.1128/MMBR.00031-15. PMID: 26864431; PMCID: PMC4771367.

92. Yang C, Wang X. Lysosome biogenesis: Regulation and functions. J Cell Biol. 2021 Jun 7;220(6):e202102001. doi: 10.1083/jcb.202102001. Epub 2021 May 5. PMID: 33950241; PMCID: PMC8105738.

93. Young C, Walzl G, Du Plessis N. Therapeutic host-directed strategies to improve outcome in tuberculosis. Mucosal Immunol. 2020 Mar;13(2):190–204. doi: 10.1038/s41385-019-0226-5. Epub 2019 Nov 26. PMID: 31772320; PMCID: PMC7039813.

94. Yuk JM, Yoshimori T, Jo EK. Autophagy and bacterial infectious diseases. Exp Mol Med. 2012 Feb 29;44(2):99-108. doi: 10.3858/emm.2012.44.2.032. PMID: 22257885; PMCID: PMC3296818.

95. Zhao D, Qiang L, Lei Z, Ge P, Lu Z, Wang Y, Zhang X, Qiang Y, Li B, Pang Y, Zhang L, Liu CH, Wang J. TRIM27 elicits protective immunity against tuberculosis by activating TFEB-mediated autophagy flux. Autophagy. 2024 Jul;20(7):1483–1504. doi: 10.1080/15548627.2024.2321831. Epub 2024 Mar 4. PMID: 38390831; PMCID: PMC11210901.

