## Supplemental Figures for "PE11 promotes intracellular persistence of *Mycobacterium tuberculosis* by inhibiting autophagy and lysosomal biogenesis by targeting the FLCN-lactate-TFEB signaling axis": Supplementary figures and tables-FF .pdf

**Supplementary Table 1: Resources and Antibodies**

| REAGENT or RESOURCE | SOURCE | CATALOGUE |
|---|---|---|
| **Antibodies** | | |
| Rabbit anti-LC3B (E7X4S) Ab | Cell signalling Technology, USA | Cat# 43566 |
| Rabbit anti-P62 Ab | St john Labs, UK | Cat# STJ113744 |
| Rabit anti-TFEB Ab | Cell signalling Technology, USA | Cat# 83010 |
| Rabit anti-FLCN Ab | Proteintech, USA | Cat# 11236-2-AP |
| Rabbit anti-β-actin Ab | Cell Signaling Technology, USA | Cat# 4970S |
| Rabbit anti-CD45 Ab | Abcam, USA | Cat# Ab10558 |
| Alexa Fluor 488-conjugated goat anti-rabbit Ab | Thermo Fisher Scientific, USA | Cat# A11034 |
| Alexa Fluor 488-conjugated goat anti-mouse Ab | Thermo Fisher Scientific, USA | Cat# A11029 |
| Alexa Fluor 594-conjugated goat anti-rabbit Ab | Thermo Fisher Scientific, USA | Cat# A11037 |
| **Bacterial and Viral strains** | | |
| *M. smegmatis strain* $mc^2{}_{155}$ | Kind gift from Dr Dipankar Chatterjee, IISC, Bangalore, India | |
| *E. coli* BL21 (DE3) | Kind gift from Dr Abhijit A. Sardesai, CDFD, Hyderabad, India | |
| **Chemicals peptides and recombinant proteins** | | |
| Rapamycin | Sigma-Aldrich, USA | Cat# 553210 |
| Bafilomycin | Sigma-Aldrich, USA | Cat# B1793 |
| Protease inhibitor cocktail | Roche, Germany | Cat# 04693159001 |
| Phosphates inhibitor cocktail | Roche, Germany | Cat# 04906845001 |
| DQ-ovalbumin™ | Thermo Fisher Scientific, USA | Cat# D12053 |
| LysoTracker™ Red | Thermo Fisher Scientific, USA | Cat# L7528 |

| Critical commercial assays | | |
|---|---|---|
| Micro BCA™ Protein Assay kit | Thermo FisherScientific, USA | Cat# 23235 |
| TNF-α cytokine | Thermo FisherScientific, USA | Cat# 88732476 |
| IL-1β cytokine | Thermo FisherScientific, USA | Cat# 887013A-88 |
| IL-6 cytokine | Thermo FisherScientific, USA | Cat# 88706486 |
| Iscript™ cDNA synthesis kit | Bio-RAD | Cat# 1708891 |
| TB Green™ Premix Ex Taq™ | TAKARA | Cat# RR420A |
| **Experimental models: organisms/strains** | | |
| C57Bl/6 Mice (Male/Female Any) | CDFD Animal house facility | Cat# NA |
| **Software and algorithms** | | |
| Prism 8.0 | GraphPad | Cat# NA |
| ImageJ | Open Source | Cat# NA |
| **Microscopy** | | |
| Leica sp8 microscope | Lieca | Cat# NA |
| **FACS** | | |
| BDLSRFortessa | BD Bioscience | Cat# NA |

**Supplementary Table 2: Primer list used for RT-PCR**

| Primer | Sequence (5'→3') | | |
|---|---|---|---|
| atg5_fp | AGCCAGGTGATGATTCACGG | Eurofins | Cat# NA |
| atg5_rp | GGCTGGGGGACAATGCTAA | Eurofins | Cat# NA |
| becn1_fp | AGGCGAAACCAGGAGAGAC | Eurofins | Cat# NA |
| becn1_rp | CCTCCCCGATCAGAGTGAA | Eurofins | Cat# NA |
| atp6v$_0$e_fp | GCATACCACGGCCTTACTGT | Eurofins | Cat# NA |
| atp6v$_0$e_rp | TGATAACTCCCGGTTAGGAC | Eurofins | Cat# NA |
| atp6v$_1$h_fp | GGATGCTGCTGTCCCAACTAA | Eurofins | Cat# NA |
| atp6v$_1$h_rp | TCTCTTGCTTGTCCTCGGAAC | Eurofins | Cat# NA |
| tfeb_fp | GGGTTGGAGCTGATATGTAGCA | Eurofins | Cat# NA |
| tfeb_rp | AAGGTTCGGGAGTATCTGTCTG | Eurofins | Cat# NA |
| 18s_fp | TTCCGATAACGAACGAGACTCT | Eurofins | Cat# NA |
| 18s_rp | TGGCTGAACGCCACTTGTC | Eurofins | Cat# NA |
| pe11_fp | CGATTCGATTGGGGAAACGG | Eurofins | Cat# NA |
| pe11_rp | AAACAGCATCGACGTCACGA | Eurofins | Cat# NA |
| sigA_fp | TGCAGTCGGTGCTGGACAC | Eurofins | Cat# NA |
| sigA_rp | GTCGCGCAGGACCTGTGAG | Eurofins | Cat# NA |
| β-actin_fp | TCGCTGCGCTGGTCGTC | Eurofins | Cat# Na |
| β-actin_rp | GGCCTCGTCACCCACATAGGA | Eurofins | Cat# Na |

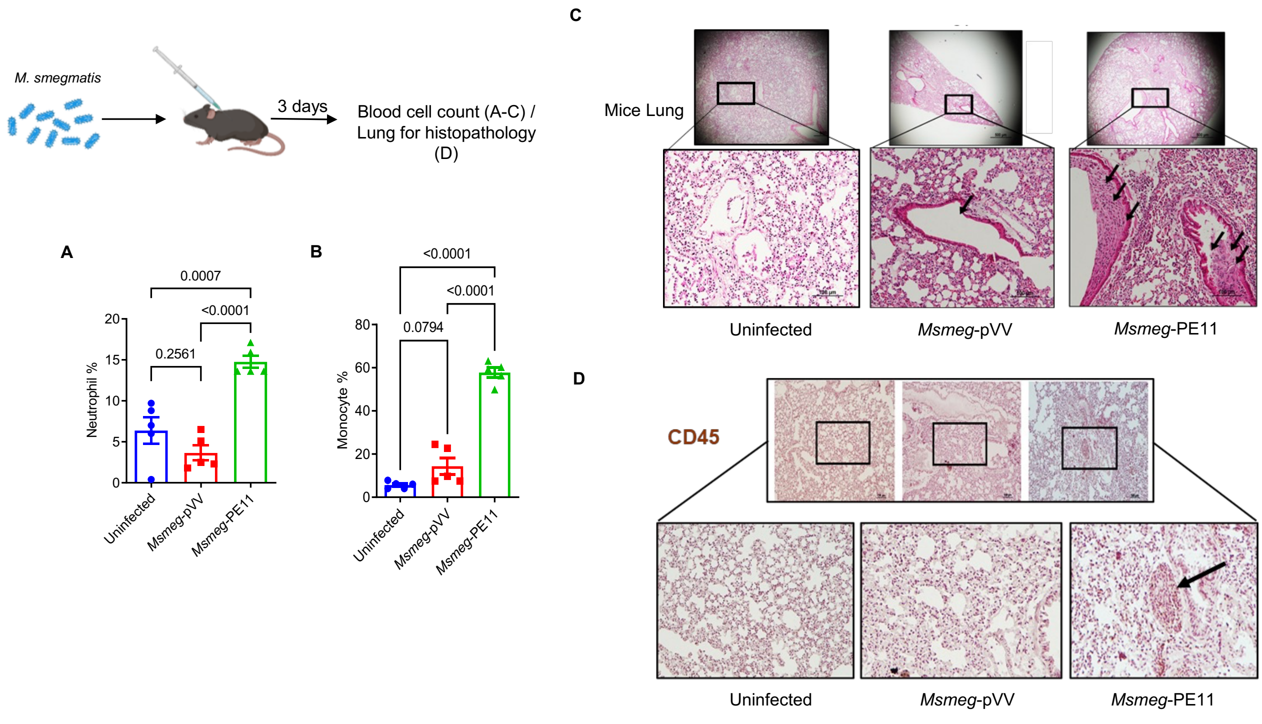

**Supplemental Figure 1.** Mice infected with M. smegmatis expressing Mtb PE11, exhibit neutrophilia, monocytosis and increased infiltration of CD45+ immune cells into the lungs. C57BL/6 mice were infected with 50 × $10^6$ CFUs of either *Msmeg*-pVV or *Msmeg*-PE11 through the intraperitoneal route. Uninfected mice were used as healthy control. At day 3 post infection, blood samples were collected and analyzed for levels of (A) Neutrophils and (B) Monocytes. Data are representative of mean ± SEM of 5 mice. (C) Also, lungs were harvested and paraffin sections were prepared and stained with haematoxylin and eosin. Scale bar of top panel is 500 µm and bottom panel is 100 µm. (D) Lung sections were probed with anti-CD45 antibody and counterstained with haematoxylin and lung tissue from uninfected mice was taken as healthy control. Arrows indicate sign of inflammation and immune cell accumulation. Photographs of representative sections were visualized at 20X magnification. Scale bar of top lane is 100 µm and bottom lane is enlarged image of the section. Significance was calculated using one-way ANOVA

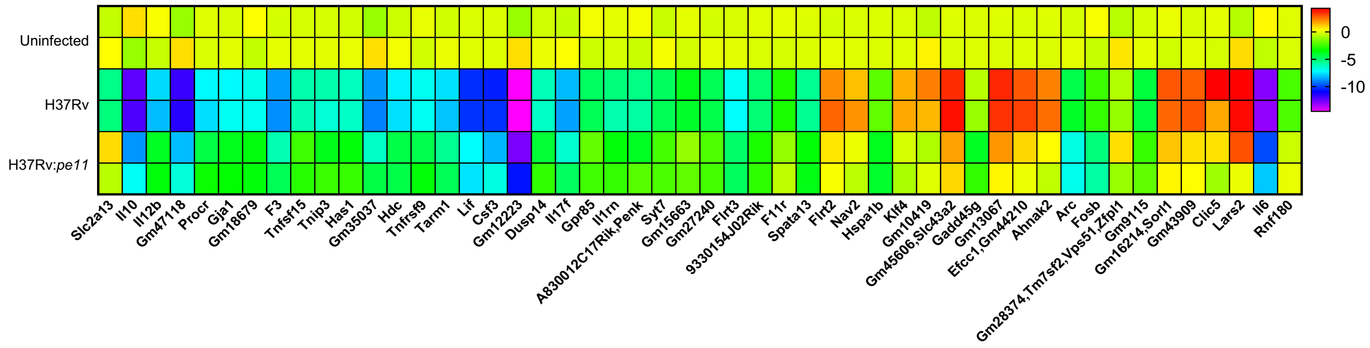

**Supplementary Figure 2. Heat map of top 50 differentially expressed genes.** C57BL/6 mice peritoneal macrophages were infected (1:5 MOI, 12 hpi), RNA sequencing was carried out. Heat-map of DEGs (columns) across infected macrophages (Rows), clustered by expression patterns is presented. Color scale: log2 normalized counts.

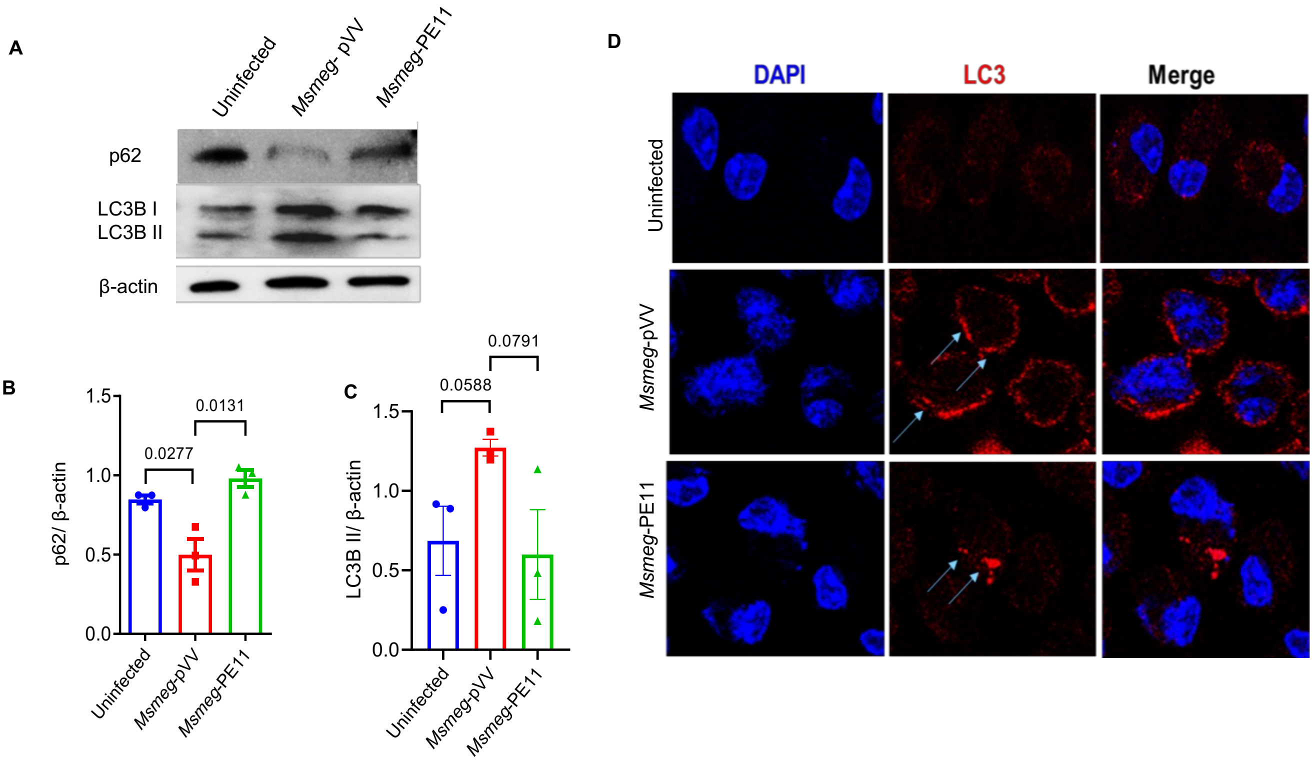

**Supplementary Figure 3. PE11 suppresses autophagy and dampened lysosomal acidification during *M. smegmatis* infection.** Analysis of autophagy markers (LC3B and p62) in C57BL/6 peritoneal macrophages infected with *Msmeg-pVV* or *Msmeg-PE11* at MOI 1:10 (6 hours post-infection (hpi)). **(A)** Representative Western blots of LC3B and p62. Quantification normalized to β-actin. Data show mean ± SEM of 3 experiments. **(B)** Confocal microscopy showing LC3B puncta formation in BMC2 macrophages infected as above (LC3B: red; DAPI: blue; scale bar: 10 µm). Significance was calculated using Student's *t* test.

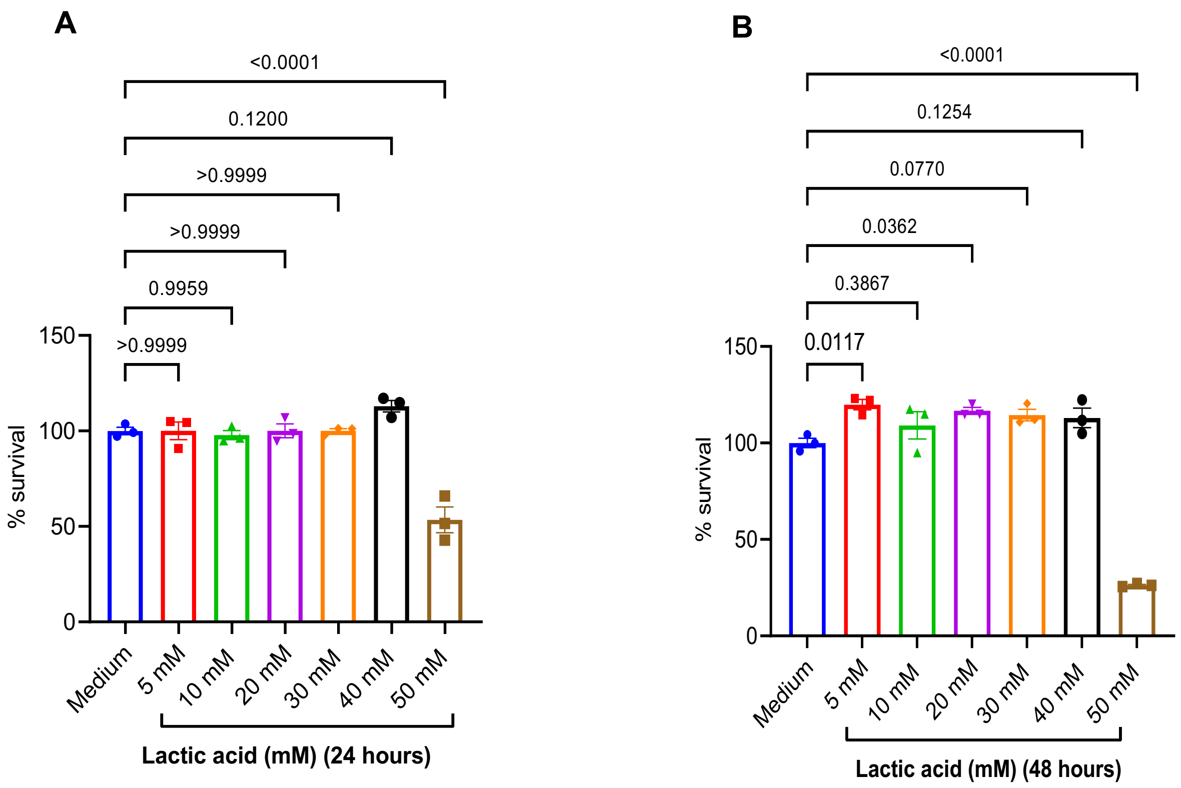

**Supplementary Figure 4. Effect of lactate supplementation on cell cytotoxicity (A, B)** Peritoneal macrophages from C57BL/6 mice were treated with various concentrations of lactate for 24 hours (A) and 48 hours (B) and cytotoxicity assay was performed by MTT assay. Data (mean ± SEM of 3 different experiments) are shown as percentage viability of the untreated cultures. Significance was calculated using one-way ANOVA

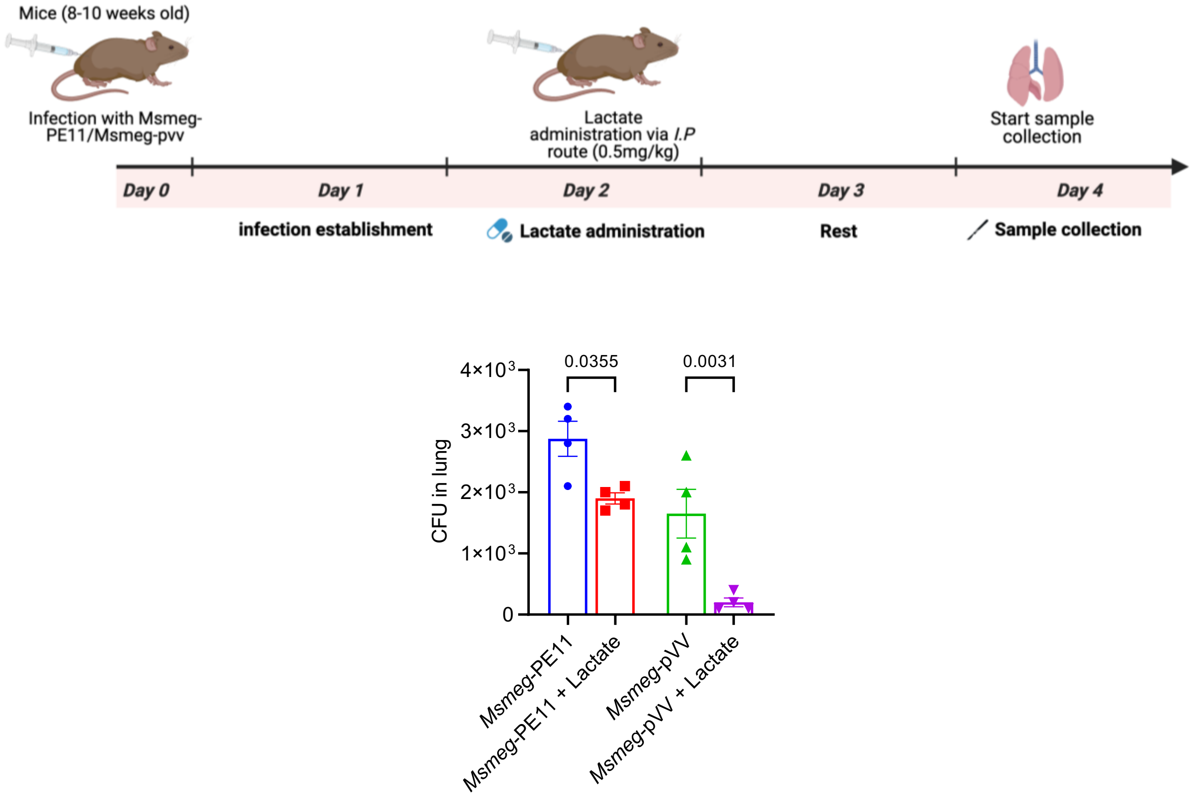

**Supplementary Figure 5. Lactic acid inhibits mycobacterial growth in mice. (A)** C57Bl/6 peritoneal macrophages infected with H37Rv (MOI 5, 12 hpi) and representative images of live macrophages with or without lactate (10 mM) are represented in the panel. Statistical significance was calculated using one-way ANOVA.
